# 3D Images of Neuronal Adhesion Molecule Contactin-2 Reveal an Unanticipated Two-State Architecture

**DOI:** 10.1101/386102

**Authors:** Z. Lu, D. Lei, S. Seshadrinathan, A. Szwed, J. Liu, J. Liu, G. Rudenko, G. Ren

**Keywords:** CNTN2, electron tomography (ET), individual-particle electron tomography (IPET), protein structure, single-molecule 3D structure, TAG-1, transmission electron microscopy (TEM)

## Abstract

Contactins (CNTNs) are important cell adhesion molecules that mediate neuronal and axoglial contacts, and lesions in these molecules are linked to neuropsychiatric disorders. The extracellular domain of CNTNs contains six Ig domains and four FNIII domains. Crystal structures have shown that Ig1-Ig4 forms a horseshoe-shaped headpiece, in which the N-terminal domains might fold back on the C-terminal domains to form molecular super-U shaped architecture. The arrangement of these domains has been controversial, which may due to the structural dynamics and conformation heterogeneity of the protein. Here, we used a single-molecule 3D imaging method, individual-particle electron tomography (IPET), to study the extracellular domain of CNTN2 that forms monomers with a broad spectrum of conformations, and obtained 60 three-dimensional (3D) reconstructions. In addition to the known horseshoe-shaped headpiece, ~75% headpieces unexpectedly adopt an open (elongated) or a semi-open conformations contributed to our understanding about structural dynamics. The ectodomains formed curve but not double-back in any uniform way, with an averaged molecular dimension of ~255 Å. The first-time demonstration of the dynamic nature and conformational preferences of the full-length CNTN2 ectodomain suggest that the headpiece exists in equilibrium in the ‘closed’ or ‘not-closed’ states. The important architecture may provide a structural platform for protein partners to influence this balance regulating the function of CNTN2. Encoding the ability of this neural adhesion molecule to form both homomers with itself, as well as recruit different protein partner to neuronal and axoglial contact points play the key role in mediating cell-cell interactions.

## INTRODUCTION

Accurate neuronal and axoglial connections are essential for the development and functioning of the nervous system. Cell surface and adhesion molecules play a crucial role in forming these connections. The cell adhesion molecules, contactins (CNTNs), form a subgroup of cell adhesion molecules that belong to ‘the immunoglobulin-like domain containing’ family. Lesions in the genes encoding CNTNs are linked to mental retardation and neuropsychiatric disorders such as autism spectrum disorder, schizophrenia and bipolar disorder ^1–3^. Since their identification in the late 1980s, CNTNs have been studied extensively, and their role in axon guidance and the formation of axoglial connections during development elucidated ^1,4–10^. Through their ability to bind different protein partners with their extracellular domain, for instance members of the CNTNAP/Caspr family and other cell adhesion molecules, CNTNs are thought to play a crucial role in the formation, organization and maintenance of neuronal and axoglial contacts, promoting the proper functioning of neuronal circuits ^11^.

All CNTNs consist of six Ig-like domains connected to four fibronectin III-like domains; the ectodomain is attached to the cell membrane by a glycosylphosphatidylinositol (GPI) anchor ^11^. Crystal structures have revealed that the first four Ig domains of CNTN2 form a compact moiety arranged in a ‘horseshoe’-shaped structure ^12,13^, a configuration that is also seen in the Ig1-Ig4 fragment of CNTN3, CNTN4, CNTN5 and CNTN6 ^14,15^. To form the U-shaped arrangement, Ig1 reaches across to interact with Ig4, while Ig2 binds to Ig3 and is connected also via a six residue linker. This horseshoe-shaped configuration is also observed in other proteins containing a concatenation of 4 or more Ig domains, such as the cell adhesion molecules neurofascin, DSCAM, DCC, and the sidekicks, as well as in hemolin, a protein involved in insect immunity ^16–21^. Crystallographic studies on other fragments have revealed that CNTN3 Ig5-FNIII_2 adopts an extended conformation while FNIII_1-FNIII_3 in CNTN1 through CNTN6 adopts an L-shaped conformation with a kink between FNIII_2 and FNIII_3 ^15^.

The CNTN Ig1-Ig4 headpiece is important functionally because it binds to protein partners. Ig1-Ig4 of CNTN2 binds NgCAM (a six Ig and five FNIII domain containing neural cell surface adhesion molecule) ^22^, while the Ig1-Ig4 regions of CNTN3 through CNTN6 (but not CNTN1 or CNTN2) bind the carbonic anhydrase domain of PTPRG ^14,15^. and that of DCC binds to draxin ^23^. The Ig1-Ig4 headpiece is also thought to be important for homophilic binding between CNTN2 molecules because mutations in this region are disruptive in cell aggregation and binding studies ^12,22^. Several models have been proposed delineating how the horseshoe-shaped headpiece in CNTN2, as well as those found in related molecules, mediate homophilic interaction, in particular the formation of trans-complexes spanning the space between two cells ^12,13,17,18,21^. The Ig1-Ig4 headpiece is not the only protein interaction site of CNTNs though, because FNIII domains are thought to play a role in homodimerization as well, facilitating side-by-side interactions within the same membrane surface to promote cis-complexes ^24,25^.

The three-dimensional (3D) architecture of the full-length CNTN2 is controversial. Mutational studies and negative-staining electron microscopy (NS-EM) have suggested that CNTN2 not only adopts a horseshoe-shaped Ig1-Ig4 headpiece like in the crystal structures but that the N-terminal half (Ig1-Ig6) also folds back on the C-terminal half (FNIII_1-FNIII_4) forming a compact ‘super-horseshoe’ ^22^. However, it has become increasingly clear that the structural results depend on the imaging method and the exact protein used. For example, NS-EM analysis of L1, a related protein with six Ig and five FNIII domains, suggested that while most molecules do indeed form a U-shaped Ig1-Ig4 headpiece, the rest of the molecule is elongated ^26^. By rotary-shadowing EM, however, the Ig1-Ig4 headpiece of L1 is observed as an elongated chain of Ig domains, not a horseshoe, unlike its counterpart in hemolin which is consistently U-shaped ^26^. Likewise, the five Ig domains in ICAM form a linear rod by rotary shadowing EM ^27^, while in DSCAM, the crystal structure of the N-terminal Ig1-Ig8 fragment reveals a horseshoe-shaped Ig1-Ig4 domain structure that is doubled back to contact Ig8 ^20^. DCC may also adopt a folded-back globular conformation, because mutations that disrupt the horseshoe-shaped headpiece disrupt netrin-1 mediated axon guidance, a ligand that interacts with FNIII_4 and FNIII_5 not Ig1-Ig4 ^16^. Thus, it remains unknown what the exact arrangement of CNTN2 is in 3D, if the N-terminal portion of CNTN2 folds back on its C-terminal portion, if CNTN2 contains a horseshoe-shaped Ig1-Ig4 headpiece or not and whether all CNTN2 molecules adopt a uniform conformation.

To examine the architecture of CNTN2, we embarked on structural studies. It is challenging to study the conformational flexibility and dynamics of multi-domain, flexible macromolecules using current traditional structure determination methods. X-ray crystallography typically reveals a single conformation. Single particle 3D EM reconstructions involve averaging thousands to millions of protein particle images grouped to share a single or limited view or conformation. As a result, the densities from the protein flexible portions are averaged out, and information on the exact nature of the conformation variability is lost. NMR, which is highly suited to visualize protein flexibility, is limited by the size of the proteins it can resolve. To study the structure and flexibility of highly dynamic and flexible proteins such as CNTN2, a panel of many different, independent protein particles would need to be analyzed ^28^. For this reason, we turned to a recently developed method, individual-particle electron tomography (IPET), which allows one to achieve a 3D density map at ~2 nm resolution on an individual particle from a series of tilted viewing images by electron tomography (ET) ^29,30^. IPET does not require a pre-given initial model, class averaging of multiple molecules, or an extended ordered lattice, and it is able to tolerate small tilt-errors. By collecting “snapshot” 3D structures from many different individual particles and comparing them, it is possible to analyze the range of protein structural flexibility and fluctuation within a population of protein molecules. Therefore, IPET is ideally suited to study molecules suspected to contain multiple states or a high degree of intrinsic conformational variability such as multi-domain neuronal adhesion molecules. We employed the IPET method combined with NS-EM to study the structure of the full-length human CNTN2. We demonstrate that CNTN2 adopts overall an extended conformation. Importantly, the CNTN2 Ig1-Ig4 headpiece is found not only in a closed U-shape configuration as seen in the crystal structures, but in vast majority of the particles (~75%) the headpiece is in fact open to varying extents. Our results suggest that the function of CNTNs may be controlled by opening and closing the horseshoe-shaped headpiece Ig1-Ig4, perhaps influenced by protein partners binding and shifting this equilibrium. This has fundamental implications for how CNTNs integrate into protein interaction networks at neuronal and axoglial connections, and mediate cell adhesion, neural cell migration and axon guidance.

## RESULTS

To assess the 3D architecture of the extracellular domain of human CNTN2 containing 10 domains, *i.e.*, the N-terminal six Ig domains linked to four C-terminal FNIII domains, we used NS-EM. NS-EM is ideally suited because the relatively small molecular mass of CNTN2 (~109 kDa) makes it challenging to image by cryo-electron microscopy (cryo-EM). Survey micrographs (**Fig. 1A**) and selected particles (**Fig. 1B** and **Fig. 1C**) of the CNTN2 sample showed an elongated structure like a chain. Particles segregated into two categories. Category 1 particles contained a lasso-shaped terminus (**Fig. 1B**). This lasso shape resembles the horseshoe-shaped configuration of the CNTN2 Ig1-Ig4 headpiece observed in crystal structures ^12,13^. Unexpectedly, however, many particles formed a completely elongated concatenation of domains, lacking a lasso-shaped terminus or exhibiting a half-opened configuration. Fully elongated particles were assigned to Category 2 (**Fig. 1C**). Taking a randomly selected collection of 514 particles, only ~24% of the particles fell into the horseshoe-shaped Category 1 particles, and ~25% of the particles fell into the elongated Category 2 particles, while the remaining ~51% were semi-closed and did not classify strictly to either Category 1 or 2.

**Fig. 1.**
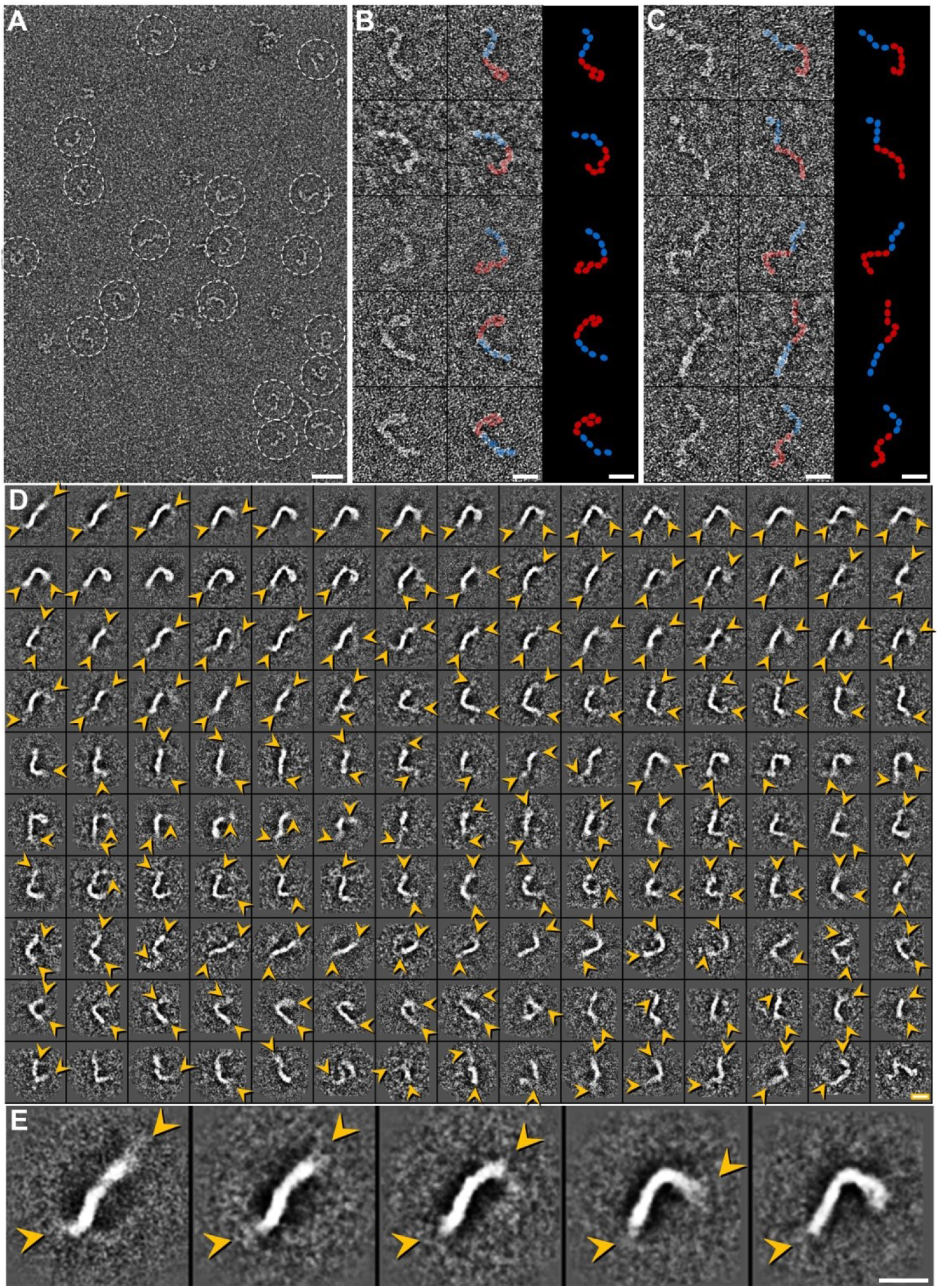
NS-EM images and reference-free 2D class averages of CNTN2. **(A)** Survey view of CNTN2 (dashed circles). **(B)** Five representative raw images of CNTN2 molecules corresponding to Category 1 particles containing a horseshoe-shaped headpiece. A schematic displays Ig (red) and FNIII (blue) domains. **(C)** Five additional representative raw images of CNTN2 particles corresponding to Category 2 particles containing a fully elongated conformation. Schematic colored as in B. **(D)** A total of 4,275 CNTN2 particles were windowed, selected, and submitted for reference-free 2D class averaging. The first 150 averages from a total of 200 classes are displayed. The number of particles in each class ranges from 6 to 96. Yellow arrows indicate blurry termini. **(E)** Zoomed-in images of five representative class averages; blurry termini are indicated by arrows. Scale bar = 400 Å for A and 100 Å for B, C, D and E.

To enhance the image contrast and reduce the noise, reference-free 2D class averaging of 4,275 particles was performed. Two hundred reference-free class averages were generated to confirm the architecture of the Category 1 (horseshoe-containing) and Category 2 (extended) particles. However, most of the class averages showed that one or both distal ends of the particles were blurry compared to the raw images (**Fig. 1D** and **Fig. 1E**). Furthermore, very few class averages were able to retain a signal for a horseshoe-shaped terminus, in contrast to particles in the raw images. Moreover, the particle lengths derived from the class averages were significantly shorter (on average ~180 Å) compared to those derived from raw particles (discussed further below). This indicated to us that the density for domains located at the N- and/or C-termini of the protein molecules was being flattening during the class averaging process. In addition, evidence of boundaries between individual domains that was clearly seen in the raw particles, was lacking in more than half of the class averages yielding molecules that appeared like continuous tubes. These results indicated several confounding issues at play which hindered the use of conventional 2D class averaging: i) heterogeneity along the length of the whole molecule likely due to intrinsic conformational flexibility, and ii) the inability of the class averaging procedure to sufficiently discriminate between discrete and unique conformational states of CNTN2 and identify them as separate classes. Thus, the smeared domain boundaries within the molecules and blurred termini suggested that CNTN2 was not suitable for conventional 2D class averaging and single-particle 3D reconstruction.

To characterize the 3D architecture of the extracellular domain of CNTN2, we therefore used a two pronged approach. To assess the dimensions of the CNTN2 ectodomain, we first used raw images to measure the size of CNTN2 particles (the distance from one end of the molecule to the other in the longest direction) to enable a statistical analysis of the size distribution within a large population of particles (**Fig. 2A**). Analysis of 514 particles yielded a minimum diameter of ~169 Å and a maximum diameter of ~342 Å. The majority of the molecules (~54%) ranged from 230 Å to 290 Å, with the most common diameter length being 255 ± 5 Å as obtained by Gaussian fitting. For comparison, we analyzed the longest dimension of 155 CNTN2 molecules from 200 reference-free class averages demonstrating that the majority of the molecules (~75%) ranged from 150 - 210 Å; the most common diameter length was 205 ± 5 Å as obtained by Gaussian fitting (**Fig. 2B**). Theoretically, the largest length of a Category 1 CNTN2 molecule containing a horseshoe-shaped headpiece would be ~340 Å, which comprises the 94 Å long Ig1-Ig4 headpiece, the ~165 Å Ig5-FNIII_2 fragment, and the ~85 Å FNIII_3-FNIII_4 fragment (based on the crystal structures PDBID:1CS6, 5I99 and 5E7L). The largest theoretical length of an elongated Category 2 CNTN2 molecule containing a fully open headpiece would be ~430 Å, i.e. 90 Å longer, because of the ~80 Å extended N-terminal Ig1-Ig4 headpiece. From this analysis of raw particles, we conclude that the imaged CNTN2 particles have a relatively wide range of diameters. Most of the CNTN2 particles are curved to some extent, in addition to the state of their Ig1-Ig4 headpiece, so that their largest dimension is not a pure summation of the individual domains. Indeed, fully extended molecules (> 320 Å; 5% of the total) as well as completely doubled-back molecules, *i.e*. the so-called super U (< 200 Å; 5% of the total) appear equally rare. In addition, we conclude that for neural adhesion molecules like CNTNs, it is critical to assess a panel of individual raw particles to obtain unbiased dimensions, because particles from the class-averages are rendered artificially shorter.

**Fig. 2.**
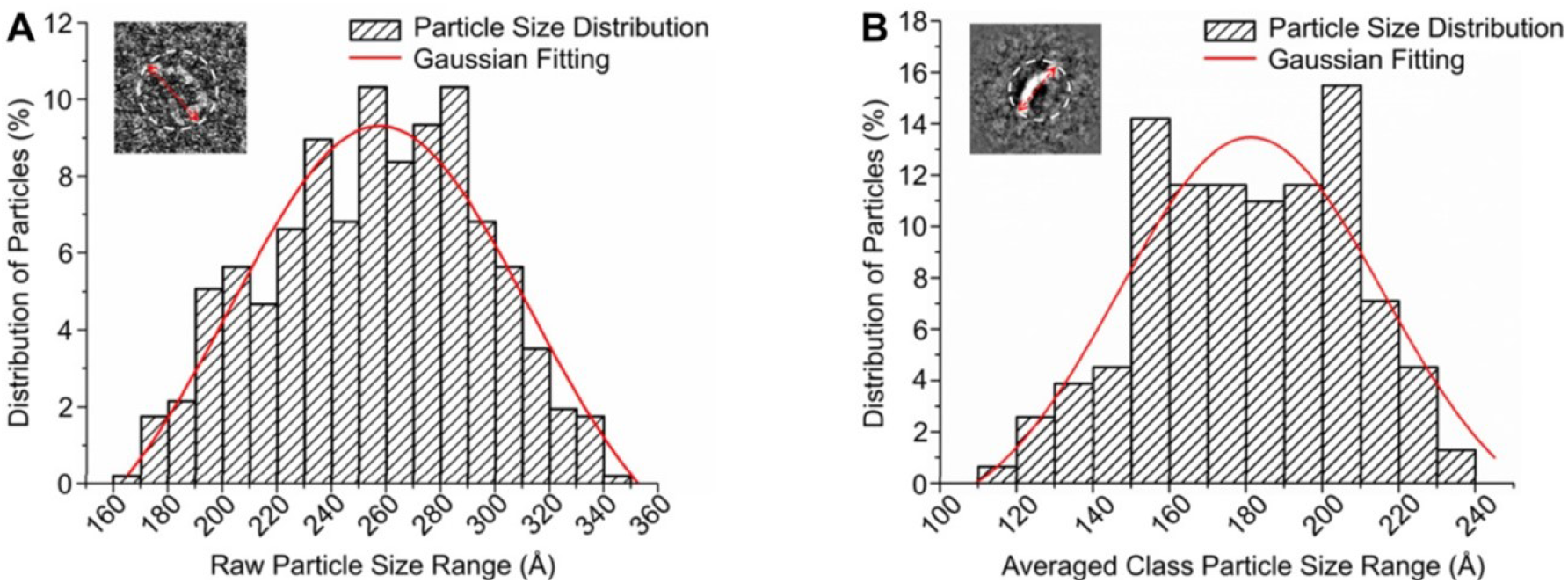
Histogram of the CNTN2 diameters. CNTN2 particles were measured in their longest dimension (red double-headed arrow) based on **(A)** raw particles and **(B)** reference-free class averages. The particle dimensions are displayed in a histogram and the distribution of the particle diameters fitted by a Gaussian curve to facilitate interpretation.

In our second approach, we performed 3D reconstructions for a panel of targeted CNTN2 particles using IPET to further reveal the 3D architecture of CNTN2. A total of 60 3D density maps were reconstructed by IPET with each one corresponding to a different individual CNTN2 particle (**Fig. 3** and **4**, **Figure S1** - **S28** and **Video S1**). The resolutions of the 60 3D density maps were in the range of 18.5-19.2 Å (see **Table S1** for details). Four representative 3D density maps (two horseshoe-shape containing Category 1 particles and two fully elongated Category 2 particles) are displayed in **Fig. 3**. The panel of final 3D reconstructions reveals each CNTN2 molecule as a chain of discrete globular densities corresponding to Ig and FNIII domains (**Fig. 4**). The ensemble of molecules reveals a wide range of overall conformations, in addition to a two-state Ig1-Ig4 headpiece that can be ‘open’ or ‘closed’ (**Fig. 4**).

**Fig. 3.**
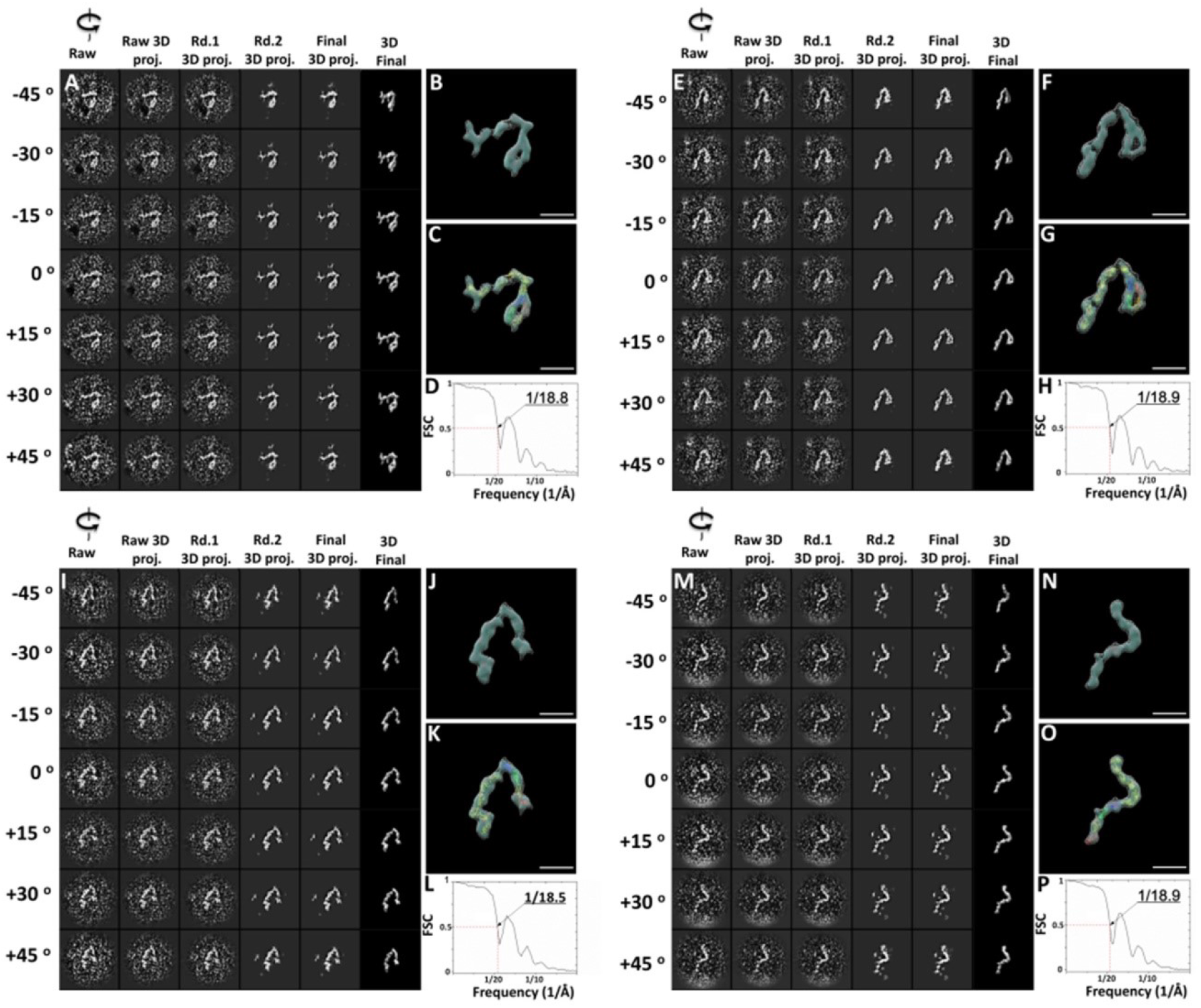
IPET 3D reconstruction of four representative CNTN2 molecules. **(A-D)**: IPET procedure for a targeted CNTN2 Category 1 particle containing a horseshoe-shaped headpiece. **(A)** Seven representative tilting views (leftmost column) of this Category 1 particle were selected from 81 tilting ET micrographs after CTF correction. The corresponding tilting projections of the 3D reconstruction from major iterations are displayed beside the raw images in next four columns. Final 3D reconstruction at the corresponding tilting angles is displayed in the rightmost column. **(B)** Final IPET 3D density map of the targeted individual particle displayed using two contour levels, *i.e*., 0.516 and 0.302. **(C)** The final IPET 3D density map with docked crystal structures, see text. **(D)** The resolution of the IPET 3D density map is 18.8 Å according to FSC analyses. **(E-H):** IPET procedure for a second targeted CNTN2 Category 1 particle. Final 3D density map is displayed using two contour levels, 0.588 and 0.305. The resolution of the IPET 3D density map is 18.9 Å according to FSC analyses; **(I-L):** IPET procedure for an individual CNTN2 Category 2 particle (elongated shape). Final 3D density map is displayed using two contour levels, 0.473 and 0.299. The resolution of the IPET 3D density map is 18.5 Å according to FSC analyses. **(M-P):** IPET procedure for a second CNTN2 Category 2 particle (elongated shape). Final 3D density map is displayed using two contour levels, 0.706 and 0.228. The resolution of the IPET 3D density map is 18.9 Å according to FSC analyses; Color coding convention: Ig1, red; Ig2, yellow; Ig3, green; Ig4, blue; and Ig5-FNIII_4, gold. Scale bar = 100 Å.

**Fig. 4.**
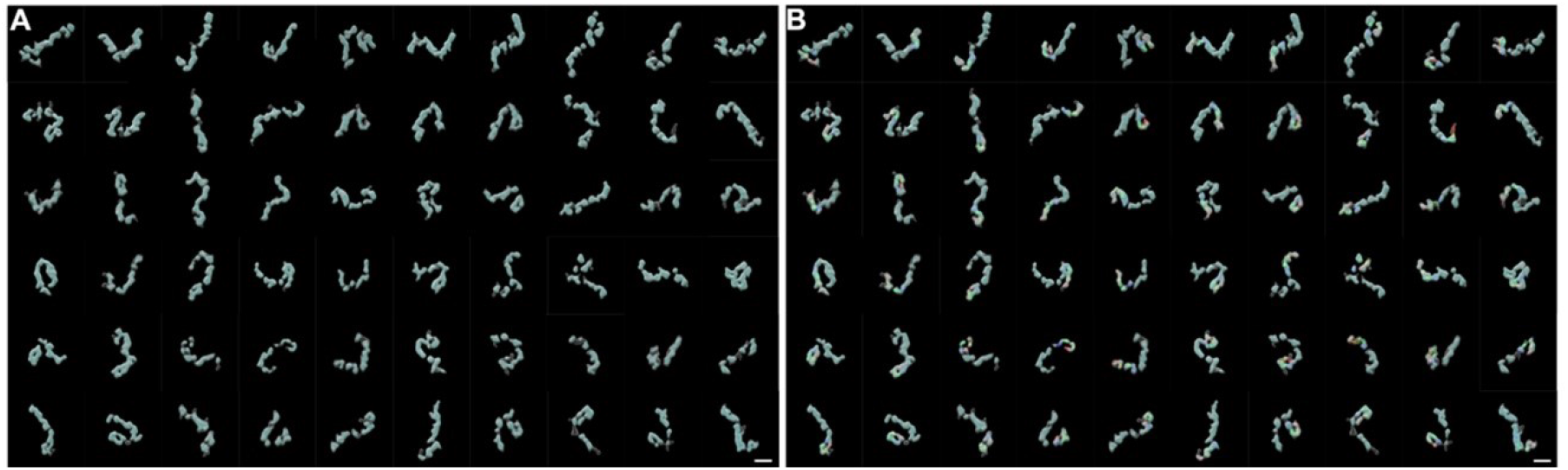
IPET 3D structures and conformational preference of 60 CNTN2 particles. **(A)** Final IPET 3D density map for 60 CNTN2 particles shown at two contour levels (see **Fig. 3**, **Fig. S1** through **S28**, and Supplementary **Table 1** for details). Inner contoured surface in light cyan and outer contour surface as silver mesh. (B) Flexibly docked fragments of CNTN2 in the IPET 3D maps. For details, see text. Color coding convention: Ig1, red; Ig2, yellow; Ig3, green; and Ig4, blue. Scale bar = 100 Å.

Our panel of IPET 3D density maps enables us to delineate the architecture of CNTN2 and its conformational preferences. To guide in the interpretation, we docked known fragments into the IPET 3D density maps. In Category 1 particles, the characteristic horseshoe-shaped Ig1-Ig4 headpiece was used to assign the N-terminal end of the molecule. By flexibly docking Ig1-Ig4, a near perfect match was obtained between the 3D envelop and the crystal structure both in terms of size and shape (**Fig. 3**). A sharp bend at the linker region between Ig2 and Ig3 was clearly observable, enabling the linear Ig1-Ig2 tandem to fold back upon the linear Ig3-Ig4 tandem (**Fig. 3A-C** and **Fig. 3E-G**). To understand the structure of entire extracellular domain, the domains of lg5 - lg6 and FNIII1 - FNIII2 were docked by the homology model of CNTN3 Ig5 – FNIII_2 fragment (PDBID: 5I99; 45.3% identity to the corresponding domains in human CNTN2), while the domain FNIII3 - FNIII4 were docked by the homology model of mouse CNTN2 FNIII_1 – FNIII_3 (PDBID: 5E7L) in each IPET 3D density maps ^15^. Similar docking studies in the IPET maps of Category 2 particles revealed a fundamentally different arrangement with the first four Ig domains in an extended conformation resembling beads-on-a-string (**Fig. 3 I-K** and **Fig. 3 M-O**). Even though the resolution of the IPET 3D density maps was insufficient to unambiguously identify the N- and C-terminal ends in the case of Category 2 particles, the N-terminal Ig and C-terminal FNIII domains are similar in size, and thus fragments could be easily docked. Our docking studies establish that the particles imaged accommodate no more than 6 Ig and 4 FNIII domains, and thus represent CNTN2 monomers. Of the 60 IPET 3D reconstructions, only ~27% contained a horseshoe-shaped Ig1-Ig4 configuration, while ~30% of the particles contained fully open, elongated Ig1-Ig4 headpieces; the remaining ~43% were semi-closed to varying extents (**Fig. 4**), i.e., very similar to what was observed for raw particles. We conclude that the majority of the CNTN2 molecules observed here by IPET (~75%) do not have a closed horseshoe-shaped Ig1-Ig4 headpiece.

To quantitatively investigate the architecture of the Ig1-Ig4 headpiece and its preference for adopting closed, open, and semi-opened states, we measured the distances between the N-terminus of Ig1 (nitrogen atom of Thr^4^) and the C-terminus of Ig4 (carbon atom of Ala^384^) after flexible docking in 60 IPET-3D density maps (**Fig. 5**; **Table S1**). The largest distance between Ig1 and Ig4 was ~179 Å (corresponding to a fully elongated Ig1-Ig4 arrangement), and the smallest distance was ~6 Å (corresponding to a horseshoe-shaped Ig1-Ig4 configuration). The distributions of distances revealed that while the most uniform conformation was the horseshoe-shaped headpiece, with an inter-domain distance of ~10-20 Å (~25% of the particles), the vast majority of particles in fact did not contain a closed, horseshoe-shaped Ig1-Ig4 headpiece (> ~20 Å), consistent with the analysis from the raw particles. Thus our results show that the architecture of CNTN2 adopts a very high degree of conformational freedom, and that the Ig1-Ig4 headpiece has a two-state architecture formed by equilibrium between a non-closed and a closed state.

**Fig. 5.**
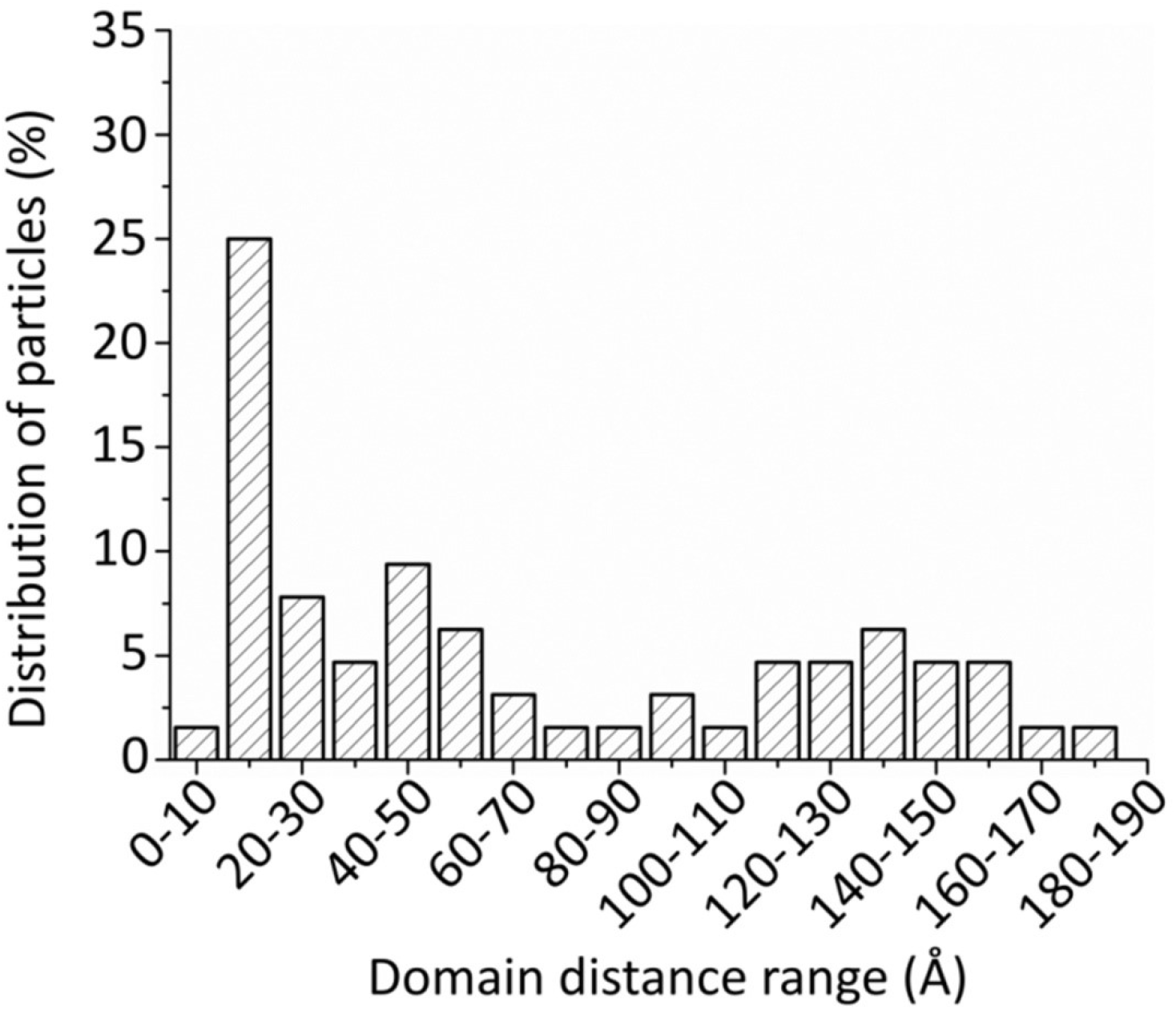
Distance distribution between Ig1 and Ig4 in 60 IPET 3D reconstructions of CNTN2. The Ig1-Ig4 headpiece (PDBID: 2OM5) was docked into IPET 3D density maps and the distance between the N-terminus of Ig1 and C-terminus of Ig4 analyzed.

## DISCUSSION

In an unprecedented single molecule approach, we demonstrate that the extracellular domain of CNTN2 forms monomers with a broad spectrum of conformations, and that the majority of the N-terminal Ig1-Ig4 headpieces assume open (elongated) or semi-open states, rather than a horseshoe-shaped configuration. The majority of the CNTN2 monomers are curved, with an average distance of ~230-290 Å between the N- and C-termini, but do not double-back in any uniform way.

In this study, CNTN2 samples were examined using NS-EM. Although cryo-EM is often used to study the structure of proteins under near-native conditions and avoids artefacts induced by fixatives and stains, we chose NS-EM because the extracellular domain of our CNTN2 protein has a calculated molecular weight of 109 kDa, which is too small to readily visualize or 3D reconstruct using cryo-EM. NS-EM typically increases the image contrast by coating protein surfaces with heavy metal salts ^31,32^. However, the conventional heavy metal staining procedure can cause artefacts, *e.g*. rouleaux formation in lipoproteins ^31,33^. For this reason, we developed an optimized NS protocol ^33^. This protocol has been validated by demonstrating that molecular envelopes obtained through our NS-EM procedure match those derived from crystal structures, for instance for the 53-kDa cholesteryl ester transfer protein ^34^, GroEL, proteasomes ^32^, as well as for flexible proteins such as an immunoglobulin (Ig)-G1 antibody and its peptide conjugates ^28,29^. NS-EM has also been applied to proteins for which no structural information was known like CNTNAP2 ^35^ and Cstn3 ^36^. Furthermore, investigation of Dscam by NS-EM using uranyl formate (as we applied here) shows that the N-terminal four Ig domains maintain a horseshoe-shaped configuration even though the rest of the molecule reveals conformational variability ^19^. Thus, proteins carrying Ig1-Ig4 headpieces can be visualized by NS-EM, and their conformations and equilibria between open and closed states assessed.

To date, all of the crystal structures of the Ig1-Ig4 headpiece in CNTNs and related proteins have revealed a compact horseshoe-shaped configuration ^12–21^. However, the data presented here reveal that most of the CNTN2 particles in our EM images in fact do not adopt a closed horseshoe-shaped Ig1-Ig4 headpiece. Our studies thus highlight the importance of examining multi-domain proteins such as CNTN2 using structural techniques that permit the identification of conformational diversity. IPET is uniquely well-suited to examine a population of individual molecules because it does not involve averaging structural information. Both EM single particle analysis and X-ray crystallography typically generate artificially uniform protein structures, because the structural information is in effect averaged, either upon processing (EM images) or intrinsically (the molecules comprising a crystal). Furthermore, crystal contacts can determine the major conformation that is observed in a crystalline environment. So although the conformation observed for CNTN2 Ig1-Ig4 in crystal structures could represent a dominant conformation, an alternative, likely explanation could be that open, flexible Ig1-Ig4 headpieces are not readily incorporating into crystals. Certainly our data suggest that an equilibrium between open (elongated) and closed (horseshoe-shaped) CNTN2 Ig1-Ig4 headpieces exists. In the related cell adhesion molecule, L1, EM studies using rotatory shadowing showed that the headpiece was open and elongated, while compact headpieces consistent with a horseshoe conformation were seen by NS-EM ^26^. In support of an equilibrium, small molecules are thought to disrupt the closed conformation of the L1 Ig1-Ig4 headpiece, altering the adhesive properties of L1 ^37,38^. Also, it has been proposed that the Ig1-Ig4 headpiece of hemolin might exist in equilibrium between extended and horseshoe-shaped conformations, and that the extended conformation would assist in mediating trans-interactions by facilitating the adhesion between opposing cell membranes ^21,26^. Interestingly, in different crystal forms of the Ig1-Ig4 headpiece of sidekick1 and sidekick 2, the horseshoe splays apart to different extents (up to 14 Å), suggesting that the crystalline environment indeed has trapped different conformations ^17^. Determining the conformational preference of CNTN2, in particular for the IG1-Ig4 headpiece is important because it has long been recognized that the dynamic properties associated with proteins are crucial determinants of their stability, folding and function ^39–41^. For example, open and closed forms of CNTN2 Ig1-Ig4 might have different adhesive properties or be preferentially recruited by partners such as CNTNAP2 ^35^.

CNTN2 functions as an adhesion molecule by forming multimeric complexes that join two cell membranes together. Our previous studies demonstrated that the CNTN2 ectodomain forms a monodisperse species by size exclusion chromatography, but its apparent molecular weight (Mw) of ~326 kDa was much larger than its calculated Mw of ~109 kDa, consistent with either an elongated or a multimeric species ^35^. In our EM images, however, we observe only monomers of CNTN2 because the molecular envelopes of these particles accommodate only six Ig and four FNIII domains. It is thus likely that CNTN2 dimers do not form at the low protein concentrations that are optimal for the preparation of NS-EM grids (0.0025 mg/ml; 2.3 nM) and/or that CNTN2 monomers have relatively low affinity for each other. Recent studies investigating the dimer formation of the related sidekick proteins determined indeed dimerization constants in the micromolar range ^17^. Dimer formation and its mechanism lie at the heart of CNTN2 function. Various very different models have been proposed for the homophilic adhesion of CNTN2 and related family members, to explain how these multimeric complexes span the extracellular space at cell contacts ^12,13,19–21,26,42^. The models differ in terms of whether CNTN2 forms cis-interactions and/or trans-interactions, how the Ig1-Ig4 headpiece mediates the interactions between CNTN2 molecules and if the FNIII domains contribute to the intermolecular interactions as well. Our studies show that CNTN2 monomers can adopt a wide range of different conformations, bending and adopting different architectures. Furthermore, given the broad range of structural variation that we observe in the Ig1-Ig4 headpiece, our data raise the possibility that the association of monomeric CNTN2 into dimers or larger oligomers involves a transition from one conformation to another, *e.g*. the closing or opening of the CNTN2 Ig1-Ig4 horseshoe.

In conclusion, by examining the individual structures of a panel of CNTN2 molecules using IPET we demonstrate that CNTN2 displays remarkable conformational variability and flexibility as well as equilibrium between an open and closed Ig1-Ig4 headpiece. These molecular properties are likely of critical importance for the ability of CNTN2 to function as a cell adhesion and cell migration guiding molecule (**Fig. 6**). Our results also demonstrate that IPET can successfully achieve meaningful intermediate resolution structures for highly flexible, conformational diverse, multi-domain proteins, yielding structural insights that are precluded by traditional EM single particle analyses and crystal structures.

**Fig. 6.**
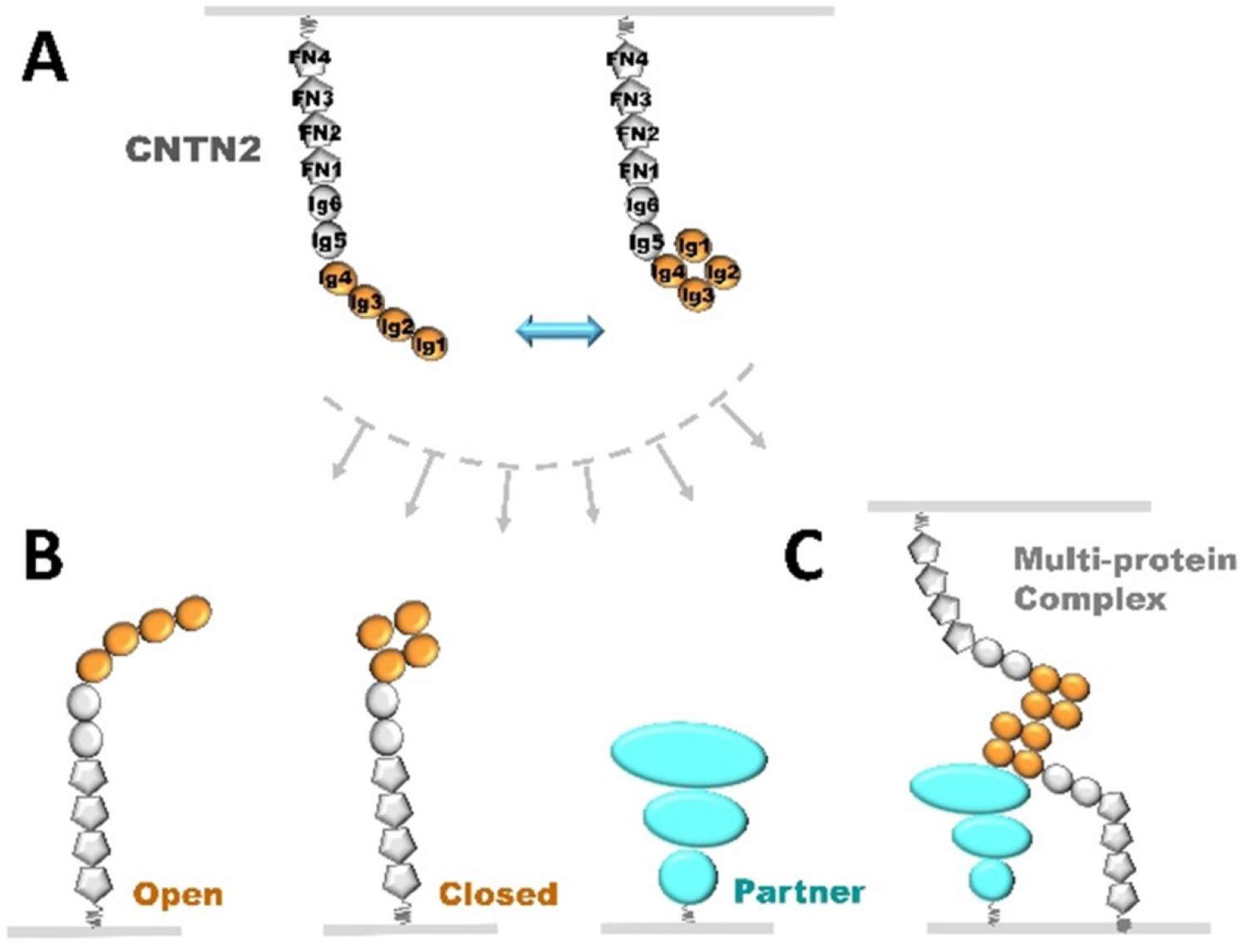
CNTN2 conformational states. **(A)** The CNTN2 ectodomain can adopt two-state architecture with its Ig1-Ig4 headpiece in equilibrium between ‘open’ and ‘closed’ conformations. **(B)** CNTN2 tethered to a neuronal membrane as in **(A)** can recruit a partner on an opposing membrane to form a trans-cellular complex, likely discriminating between CNTN2 molecules with ‘open’ or ‘closed’ Ig1-Ig4 headpieces or completely different proteins such as CNTNAP2. **(C)** The different conformational states of CNTN2 may differentially promote the assembly of complex multi-protein trans-cellular bridges, for example the CNTN2:CNTNAP2 tripartite complex at axoglial junctions.

## MATERIALS AND METHODS

### Protein expression and purification

Human contactin 2 (CNTN2) ectodomain (L^29^ESQ … VRNG^1004^) (accession: BC BC129986.1) followed by a C-terminal tag SASTSHHHHHH was produced using baculo-virus mediated overexpression in HighFive cells with Insect-XPRESS+L-Glutamine medium (Lonza). Briefly, medium containing the secreted proteins was concentrated after protease inhibitors were added, dialyzed overnight (25 mM sodium phosphate pH 8, 250 mM NaCl), and purified on a Ni-NTA column (Invitrogen, 25 mM sodium phosphate pH 8, 500 mM NaCl, eluted with an imidazole gradient). Subsequently, the protein was dialyzed into 25 mM Tris pH 8, 100 mM NaCl, applied to a MonoQ column (GE Healthcare) equilibrated with 25 mM Tris pH 8, 50 mM NaCl, and subsequently eluted with a NaCl gradient. Lastly, the sample was applied to a HiLoad Superdex-200 16/60 size exclusion column (GE Healthcare, 10 mM HEPES pH 8.0, 50 mM NaCl). Purified proteins were stored in flash-frozen aliquots.

### Negative-staining electron microscopy specimen preparation

CNTN2 samples were prepared with an optimized negative-staining (NS) method ^32,33,43^. In brief, CNTN2 at 1.0 mg/ml was diluted to 0.0025 mg/ml with 25 mM Tris pH 8, 100 mM NaCl, 3 mM CaCl_2_. An aliquot (~4 μl) of CNTN2 sample was placed on a thin-carbon-coated 200 mesh copper grid (CF200-Cu, EMS) that had been glow-discharged for 15 seconds. After ~1 min incubation, excess solution was blotted with filter paper, and the grid was stained for ~15 seconds by submersion in two drops (~35 μl) of 1% (w/v) uranyl formate (UF) on Parafilm in succession before being nitrogen-air-dried at room temperature.

### Electron microscopy data acquisition and image pre-processing

The NS micrographs were acquired at room temperature on a Gatan UltraScan 4K×4K CCD by a Zeiss Libra 120 transmission electron microscope (Carl Zeiss NTS) operating at 120 kV at 125,000× and 80,000× magnification under near Scherzer focus (0.1 μm) and defocus of 0.6 μm. Each pixel of the micrographs corresponded to 0.94 Å for 125,000× magnification and 1.48 Å for 80,000× magnification in the specimens. Micrographs were processed with EMAN, SPIDER, and FREALIGN software packages ^44–46^. The defocus and astigmatism of each micrograph were examined by fitting the contrast transfer function (CTF) parameters with its power spectrum by *ctffind3* in the FREALIGN software package ^45^. Micrographs with distinguishable drift effects were excluded and then CTF corrected with the SPIDER software ^44^. Only isolated particles from the NS-EM images were initially selected and windowed using the boxer program in EMAN and then manually adjusted. A total of 200 micrographs of CNTN2 were acquired, in which a total of 4,275 particles were windowed and selected. These particles were aligned and classified by reference-free class averaging with *refine2d.py* in the EMAN software package ^46^.

### Electron tomography data acquisition and image pre-processing

Electron tomography data of CNTN2 was acquired at 80,000× magnification under near Scherzer focus (0.1 μm) and defocus of 0.6 μm by a Gatan Ultrascan 4K × 4K pixel CCD equipped in a Zeiss Libra 120 Plus TEM operated under 120 kV. Each pixel of the micrographs corresponded to 1.48 Å for 80,000× magnification in the specimens. The specimens mounted on a Gatan high-tilt room-temperature holder were tilted at angles ranging from −60° to 60° in steps of 1.5°. The total electron dose was ~200 e^-^/Å^2^. The tilt series of tomographic data were controlled and imaged by Gatan tomography software that was preinstalled on the microscope. Tilt series were initially aligned together with the IMOD software package^47^. The defocus of each tilt micrograph was determined by *ctffind3* in the FREALIGN software package, and then CTF correction was conducted by the TOMOCTF software ^45,48^. The 3D density maps of each particles of CNTN2 was reconstructed by IPET software ^30^. A total of 17 tomograms of CNTN2 were acquired, in which, three tomograms were used for 3D reconstruction of a total of 60 particles.

### Individual-particle electron tomography (IPET) 3D reconstruction

3D density maps were reconstructed using the IPET method as described previously ^30^. Briefly, images containing only a targeted particle were selected and windowed from each tilted whole-micrograph after CTF correction. An initial model was obtained by back-projecting these particle images into a 3D map according to their tilting angles. The projections of the “initial” model were then used as references for tilting image alignment during the first iteration of refinement. The reconstruction obtained by back-projecting the aligned images were then used for generating the references for the next round of iteration. During the iterations, automatically generated Gaussian low-pass filters and masks were applied to both the references and tilt images before translational parameter searching (refining translational parameters).

### Fourier shell correlation (FSC) analysis of IPET 3D maps

The resolutions of the IPET 3D reconstructed density maps were analyzed using the Fourier shell correlation (FSC) criterion. The center-refined raw ET images were split into two groups based on having an odd- or even-numbered index in the order of tilting angles. Each group was used independently to generate its 3D reconstruction by IPET; these two IPET 3D reconstructions were then used to compute the FSC curve over their corresponding spatial frequency shells in Fourier space (using the ‘RF 3’ command in SPIDER) ^44^. The frequency at which the FSC curve falls to a value of 0.5 was used to assess the resolution of the final IPET 3D density map.

### PDB docking

The 3D density maps of CNTN2 Ig1-Ig4 domains were flexible docked by crystal structure (PDB accession code: 2OM5) using Chimera ^49^, in which, each domain of the structures was treated as a ridge body, the links between the domains were flexible in structure. The U-shape of Ig1-Ig4 was used as a marker to identify the N-terminus of the CNTN2 molecules in density maps. For maps where the Ig1-Ig4 headpiece was elongated the assignment of the N- and C-termini are ambiguous. The CNTN2 lg5, lg6 and FNIII 1 – FNIII4 domains in the 3D density maps were docked by the homology models of CNTN3 Ig5–FNIII_2 fragment (PDBID: 5I99; 45.3% identity to the corresponding domains in human CNTN2), and the mouse CNTN2 FNIII_2 – FNIII_3 (PDBID: 5E7L) ^15^.

### Statistical analysis of the CNTN2 particle size and distance between the Ig1 and Ig4 domains

For statistical analyses of the particle size and shape, a total of 514 particles selected from a total of 64 micrographs were used for measurements. For the measurement, each particle were presented by a oval particle, in which the longest and shortest diameters were used for statistical calculation. A histogram of the particle dimensions was generated with a sampling step of 10 Å. The histogram was fitted with a Gaussian function in Origin 7.5 to facilitate data analysis. For statistical analysis of the domain distance between Ig1 and Ig4, a total of 60 IPET 3D density maps were used. The domain distance between Ig1 and Ig4 was defined as the distance between the nitrogen atom of Thr ^4^ (Ig1) and the carbon atom of Ala^384^ (Ig4), as measured in Chimera ^49^. The histogram for the domain distance was generated with a sampling step of 10.0 Å.

## ACKNOWLEDGEMENTS

This work was funded by NIMH (R01MH077303) with additional support provided by the Sealy Center for Structural Biology and Molecular Biophysics (UTMB), and the Brain and Behavior Research Foundation to G. Rudenko. Work at the Molecular Foundry was supported by the Office of Science of the U.S. Department of Energy under Contract No. DE-AC02-05CH11231. J.L. and G. Ren were supported by the National Heart, Lung, and Blood Institute of the National Institutes of Health (R01HL115153) and the National Institute of General Medical Sciences of the National Institutes of Health (R01GM104427).

## Data Deposition

3D maps have been deposited in the wwPDB, accession codes EMD-7353 through EMD-7413 for density maps in **Fig. 3** and **Figure S1** through **Fig. S28**.

---

**Figure S1.**
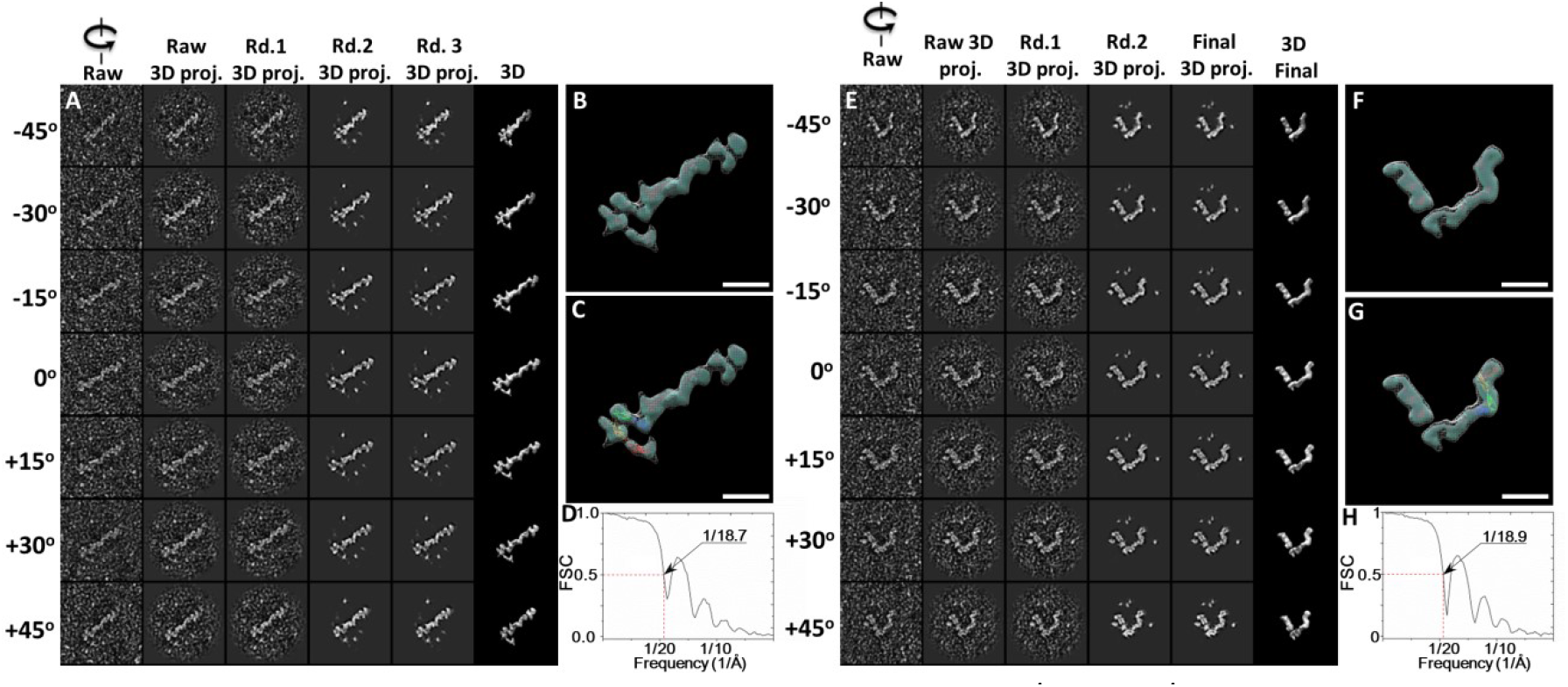
Detailed IPET 3D reconstruction process of the 5^th^ and 6^th^ CNTN2 molecules. **(A)** Seven representative tilting views (leftmost column) of the 5^th^ individual particle were selected from 81 tilting ET micrographs after CTF correction. The corresponding tilting projections of the 3D reconstruction from major iterations are displayed beside the raw images in next four columns. Final 3D reconstruction at corresponding tilting angles is displayed in the rightmost column. **(B)** Final IPET 3D density map of the targeted individual particle displayed using two contour levels, *i.e*., 0.211 and 0.125. **(C)** Final 3D density map with the docked Ig1-Ig4 crystal structure. **(D)** The resolution of the IPET 3D map is ~18.7 Å according to FSC analyses. **(E-H)** The 3D density map of the 6^th^ individual particle of CNTN2 was reconstructed from the tilt images using IPET. Final 3D density map was displayed using two contour levels, 0.326 and 0.159. The resolution of the IPET 3D map is ~18.9 Å according to FSC analyses. Scale bar = 100 Å.

**Figure S2.**
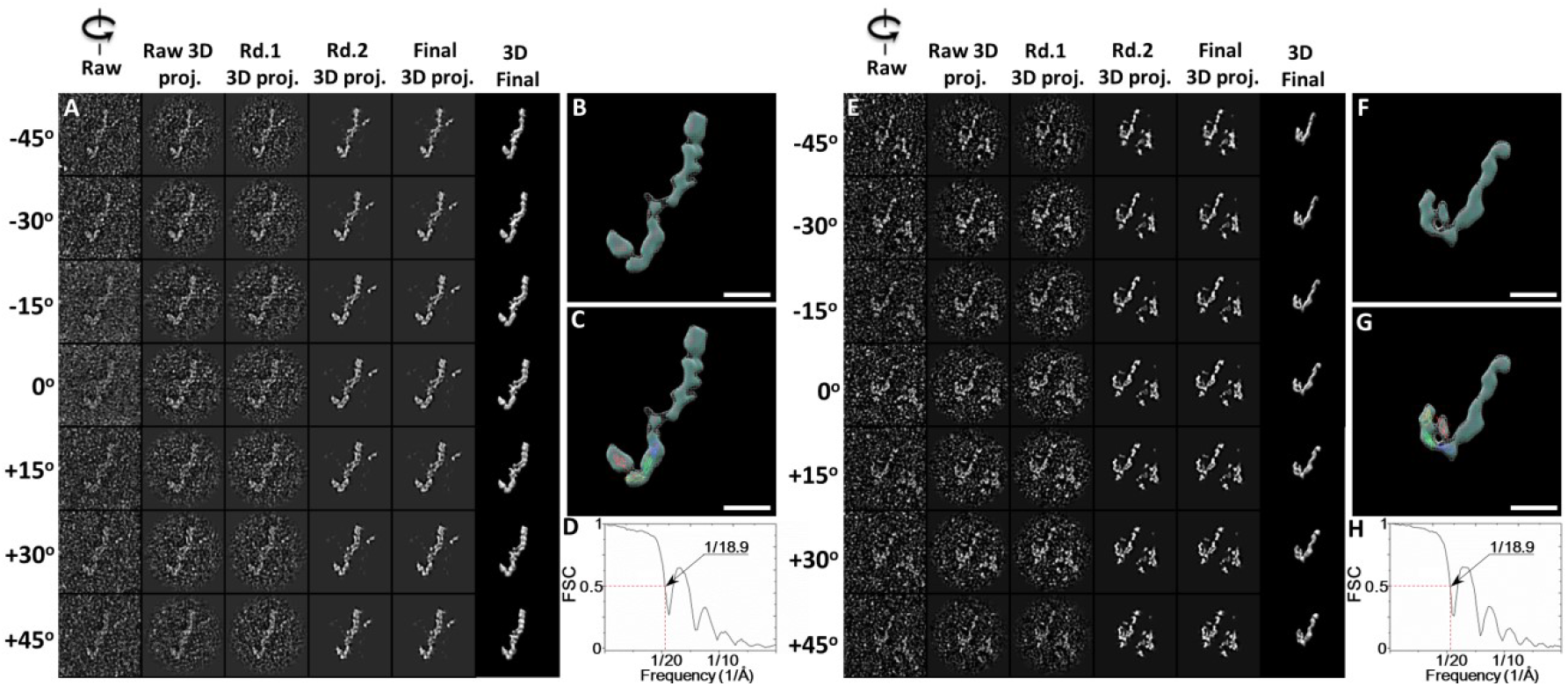
Detailed IPET 3D reconstruction process of the 7^th^ and 8^th^ CNTN2 molecules. **(A)** Seven representative tilting views (leftmost column) of the 7^th^ individual particle were selected from 81 tilting ET micrographs after CTF correction. The corresponding tilting projections of the 3D reconstruction from major iterations are displayed beside the raw images in next four columns. Final 3D reconstruction at corresponding tilting angles is displayed in the rightmost column. **(B)** Final IPET 3D density map of the targeted individual particle displayed using two contour levels, *i.e*., 0.230 and 0.151. **(C)** Final 3D density map with the docked Ig1-Ig4 crystal structure. **(D)** The resolution of the IPET 3D map is ~18.9 Å according to FSC analyses. **(E-H)** The 3D density map of the 8^th^ individual particle of CNTN2 was reconstructed from the tilt images using IPET. Final 3D density map was displayed using two contour levels, 0.572 and 0.342. The resolution of the IPET 3D map is ~18.9 Å according to FSC analyses. Scale bar = 100 Å.

**Figure S3.**
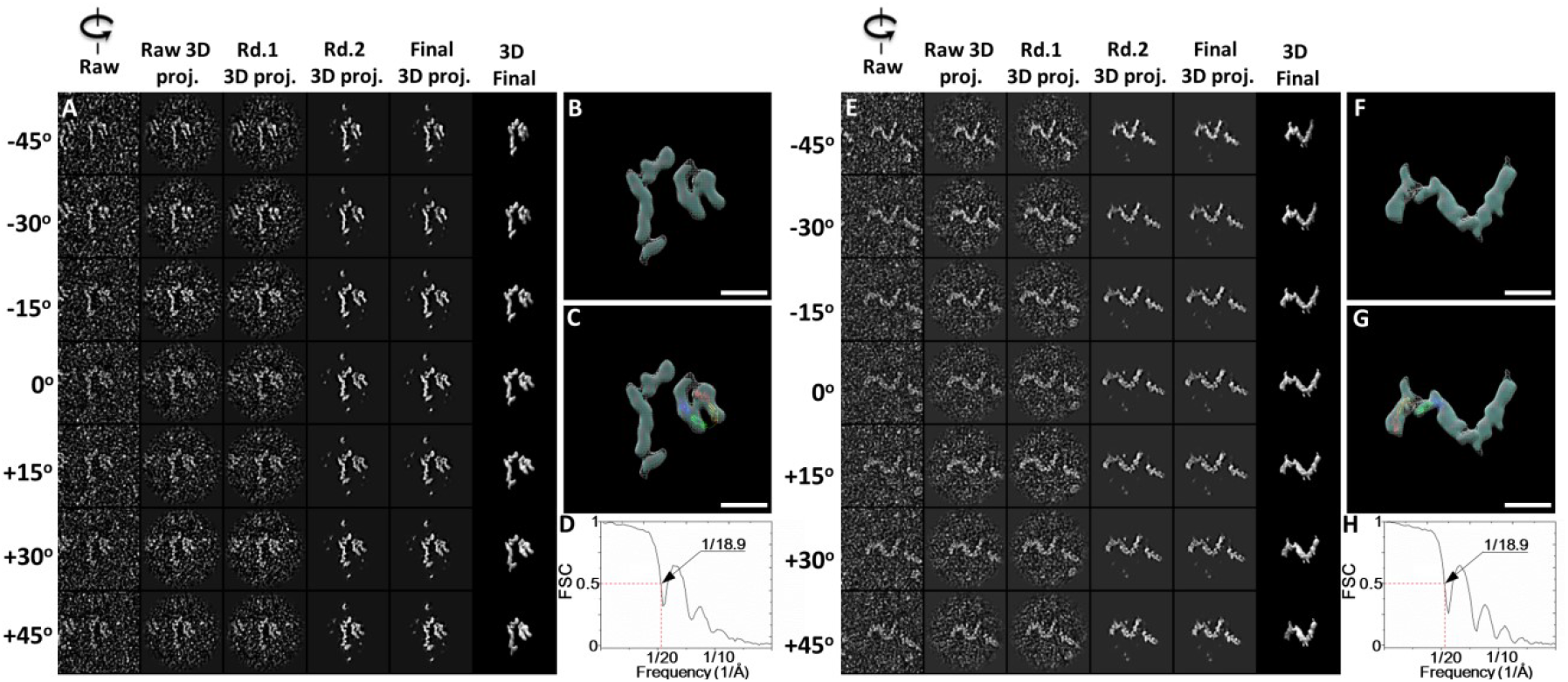
Detailed IPET 3D reconstruction process of the 9^th^ and 10^th^ CNTN2 molecules. **(A)** Seven representative tilting views (leftmost column) of the 9^th^ individual particle were selected from 81 tilting ET micrographs after CTF correction. The corresponding tilting projections of the 3D reconstruction from major iterations are displayed beside the raw images in next four columns. Final 3D reconstruction at corresponding tilting angles is displayed in the rightmost column. **(B)** Final IPET 3D density map of the targeted individual particle displayed using two contour levels, *i.e*., 0.286 and 0.169. **(C)** Final 3D density map with the docked Ig1-Ig4 crystal structure. **(D)** The resolution of the IPET 3D map is ~18.9 Å according to FSC analyses. **(E-H)** The 3D density map of the 10^th^ individual particle of CNTN2 was reconstructed from the tilt images using IPET. Final 3D density map was displayed using two contour levels, 0.421 and 0.309. The resolution of the IPET 3D map is ~18.9 Å according to FSC analyses. Scale bar = 100 Å.

**Figure S4.**
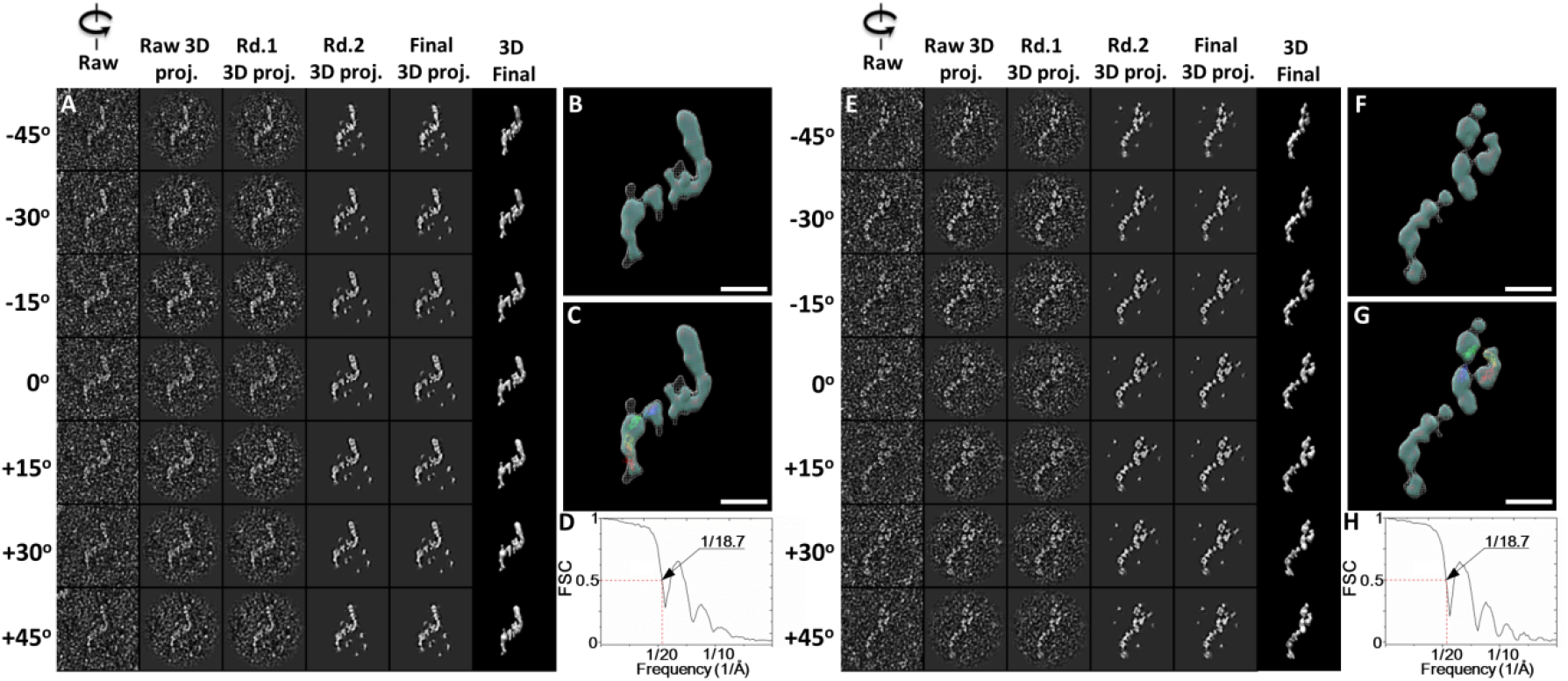
Detailed IPET 3D reconstruction process of the 11^th^ and 12^th^ CNTN2 molecules. **(A)** Seven representative tilting views (leftmost column) of the 11^th^ individual particle were selected from 81 tilting ET micrographs after CTF correction. The corresponding tilting projections of the 3D reconstruction from major iterations are displayed beside the raw images in next four columns. Final 3D reconstruction at corresponding tilting angles is displayed in the rightmost column. **(B)** Final IPET 3D density map of the targeted individual particle displayed using two contour levels, *i.e*., 0.252 and 0.135. **(C)** Final 3D density map with the docked Ig1-Ig4 crystal structure. **(D)** The resolution of the IPET 3D map is ~18.7 Å according to FSC analyses. **(E-H)** The 3D density map of the 12^th^ individual particle of CNTN2 was reconstructed from the tilt images using IPET. Final 3D density map was displayed using two contour levels, 0.267 and 0.154. The resolution of the IPET 3D map is ~18.7 Å according to FSC analyses. Scale bar = 100 Å.

**Figure S5.**
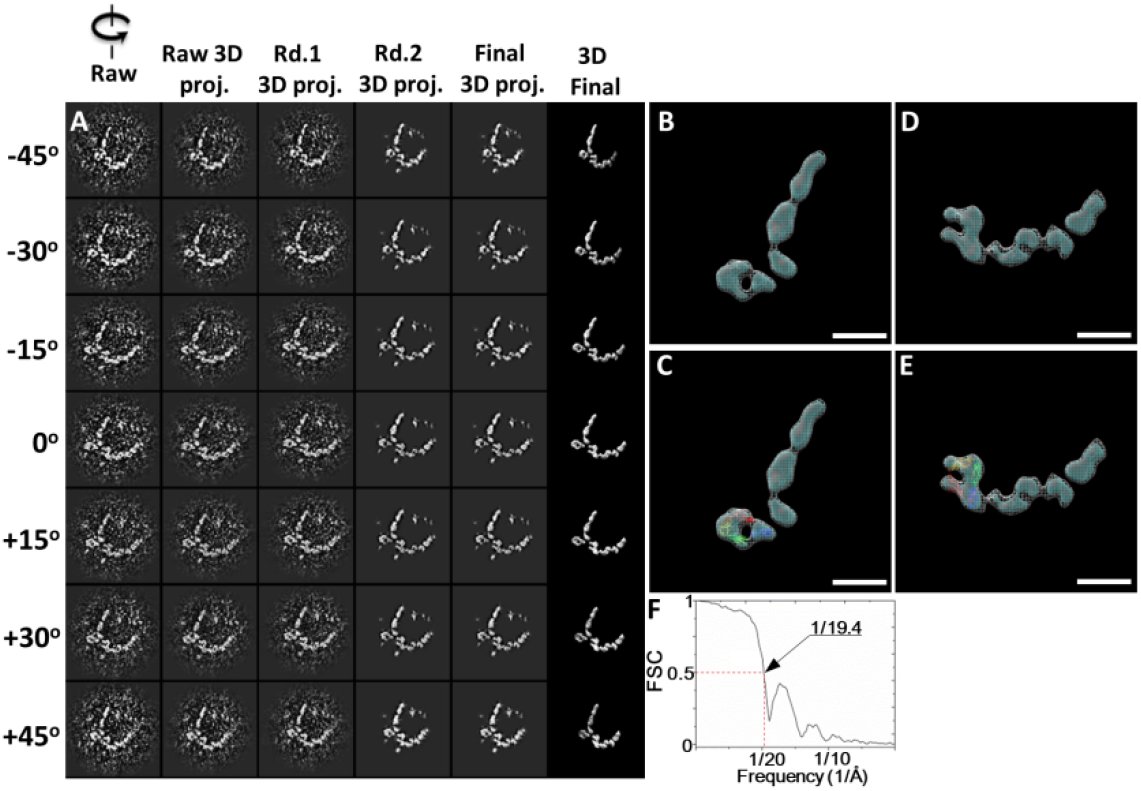
Detailed IPET 3D reconstruction process of the 13^th^ and 14^th^ CNTN2 molecule. **(A)** Seven representative tilting views (leftmost column) of the 13^th^ and 14^th^ individual particles were selected from 81 tilting ET micrographs after CTF correction. The corresponding tilting projections of the 3D reconstruction from major iterations are displayed beside the raw images in next four columns. Final 3D reconstruction at corresponding tilting angles was displayed in the rightmost column. **(B)** Final IPET 3D density map of the 13^th^ targeted individual particle displayed using two contour levels, *i.e*., 0.629 and 0.360. **(C)** Final 3D density map with the docked Ig1-4 crystal structure. **(D)** Final IPET 3D density map of the 14^th^ targeted individual particle displayed using two contour levels, *i.e*., 0.217 and 0.141. **(E)** Final 3D density map with the docked Ig1-4 crystal structure. **(F)** The resolution of the IPET 3D map is ~19.4 Å according to FSC analyses. Scale bar = 100 Å.

**Figure S6.**
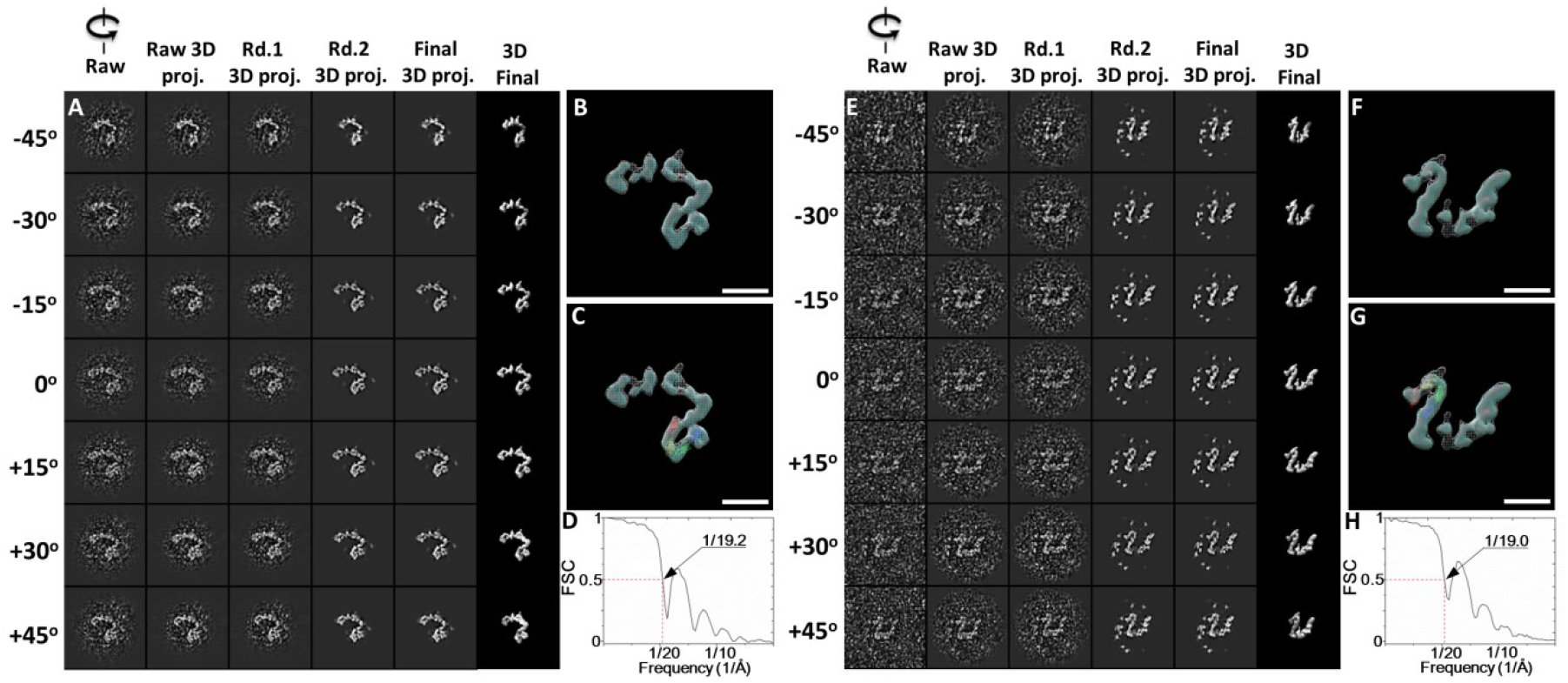
Detailed IPET 3D reconstruction process of the 15^th^ and 16^th^ CNTN2 molecules. **(A)** Seven representative tilting views (leftmost column) of the 15^th^ individual particle were selected from 81 tilting ET micrographs after CTF correction. The corresponding tilting projections of the 3D reconstruction from major iterations are displayed beside the raw images in next four columns. Final 3D reconstruction at corresponding tilting angles is displayed in the rightmost column. **(B)** Final IPET 3D density map of the targeted individual particle displayed using two contour levels, *i.e*., 0.245 and 0.151. **(C)** Final 3D density map with the docked Ig1-Ig4 crystal structure. **(D)** The resolution of the IPET 3D map is ~19.2 Å according to FSC analyses. **(E-H)** The 3D density map of the 16^th^ individual particle of CNTN2 was reconstructed from the tilt images using IPET. Final 3D density map was displayed using two contour levels, 0.529 and 0.177. The resolution of the IPET 3D map is ~19.0 Å according to FSC analyses. Scale bar = 100 Å.

**Figure S7.**
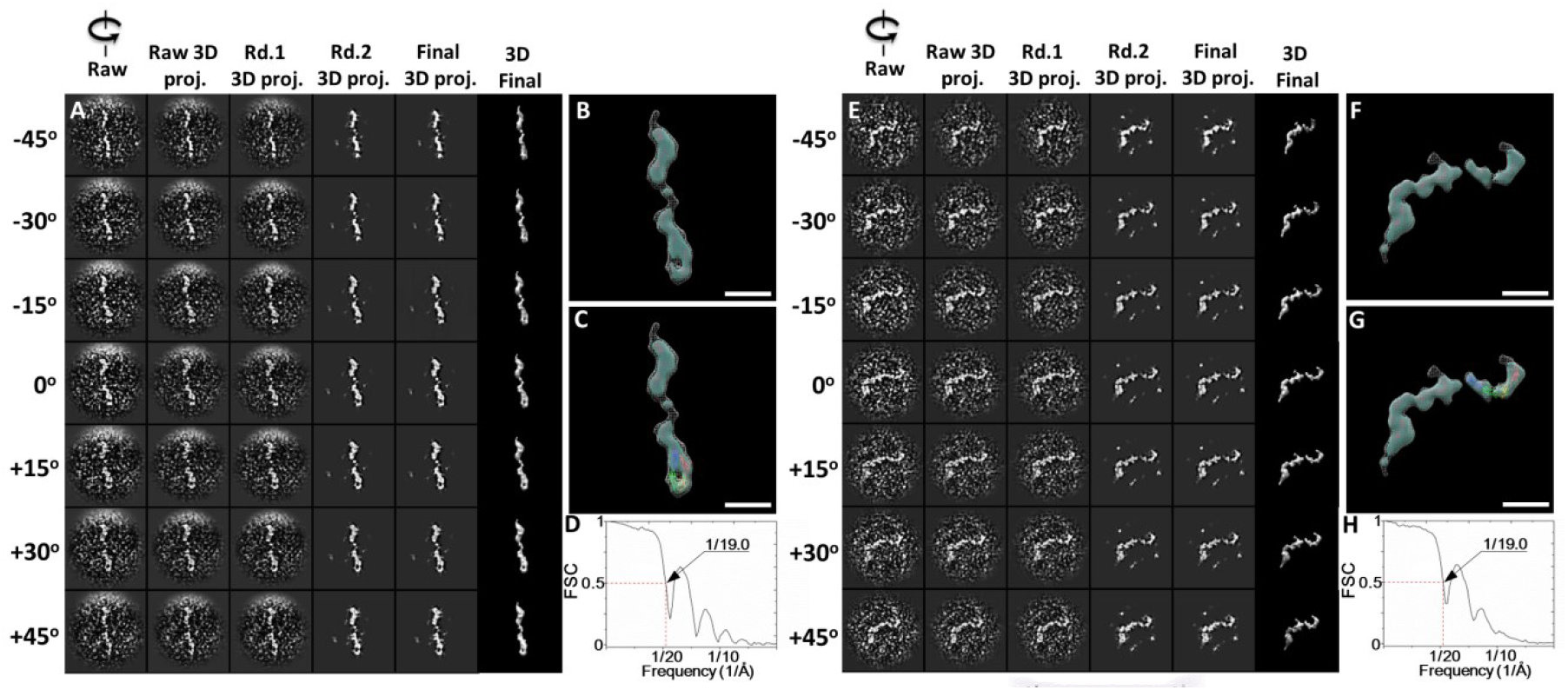
Detailed IPET 3D reconstruction process of the 17^th^ and 18^th^ CNTN2 molecules. **(A)** Seven representative tilting views (leftmost column) of the 17^th^ individual particle were selected from 81 tilting ET micrographs after CTF correction. The corresponding tilting projections of the 3D reconstruction from major iterations are displayed beside the raw images in next four columns. Final 3D reconstruction at corresponding tilting angles is displayed in the rightmost column. **(B)** Final IPET 3D density map of the targeted individual particle displayed using two contour levels, *i.e*., 0.400 and 0.258. **(C)** Final 3D density map with the docked Ig1-Ig4 crystal structure. **(D)** The resolution of the IPET 3D map is ~19.0 Å according to FSC analyses. **(E-H)** The 3D density map of the 18^th^ individual particle of CNTN2 was reconstructed from the tilt images using IPET. Final 3D density map was displayed using two contour levels, 0.668 and 0.321. The resolution of the IPET 3D map is ~19.0 Å according to FSC analyses. Scale bar = 100 Å.

**Figure S8.**
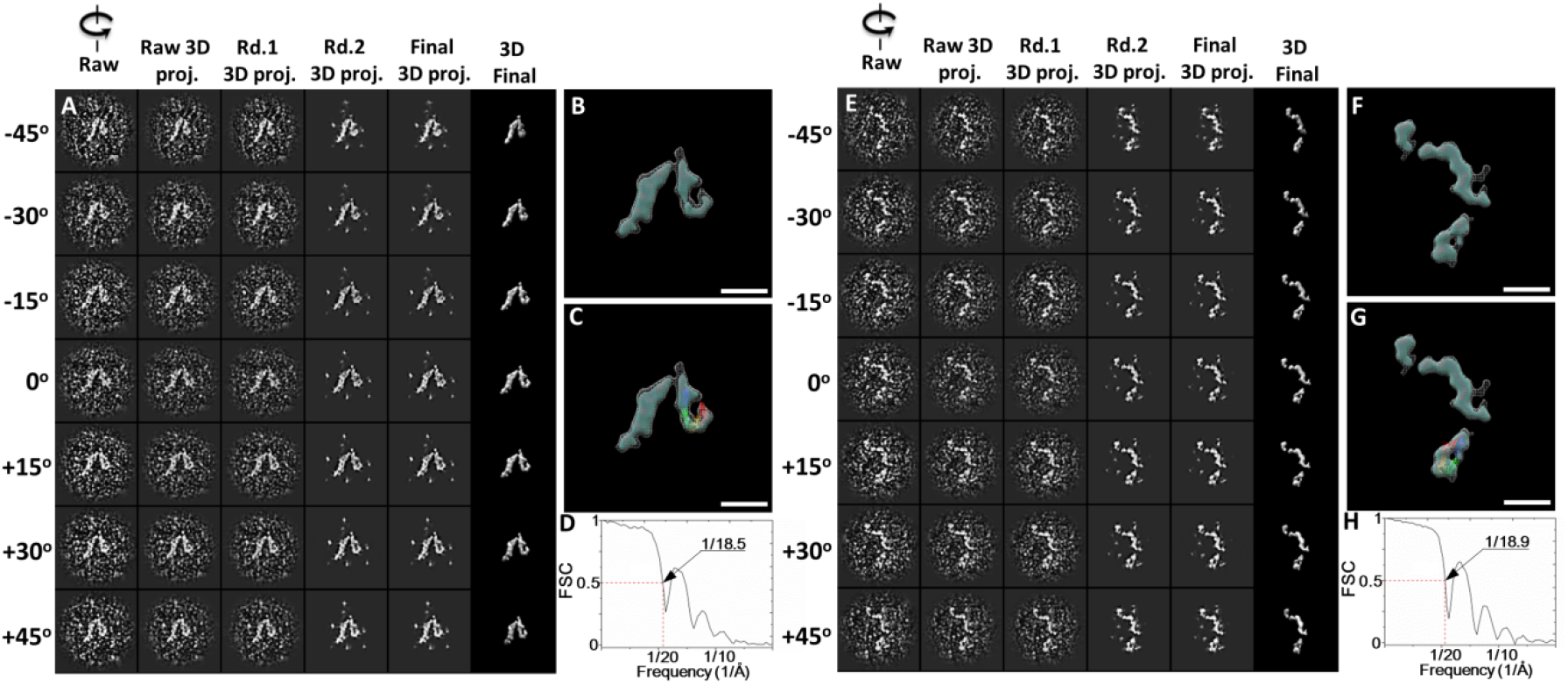
Detailed IPET 3D reconstruction process of the 19^th^ and 20^th^ CNTN2 molecules. **(A)** Seven representative tilting views (leftmost column) of the 19^th^ individual particle were selected from 81 tilting ET micrographs after CTF correction. The corresponding tilting projections of the 3D reconstruction from major iterations are displayed beside the raw images in next four columns. Final 3D reconstruction at corresponding tilting angles is displayed in the rightmost column. **(B)** Final IPET 3D density map of the targeted individual particle displayed using two contour levels, *i.e*., 0.415 and 0.236. **(C)** Final 3D density map with the docked Ig1-Ig4 crystal structure. **(D)** The resolution of the IPET 3D map is ~18.5 Å according to FSC analyses. **(E-H)** The 3D density map of the 20^th^ individual particle of CNTN2 was reconstructed from the tilt images using IPET. Final 3D density map was displayed using two contour levels, 0.355 and 0.274. The resolution of the IPET 3D map is ~18.9 Å according to FSC analyses. Scale bar = 100 Å.

**Figure S9.**
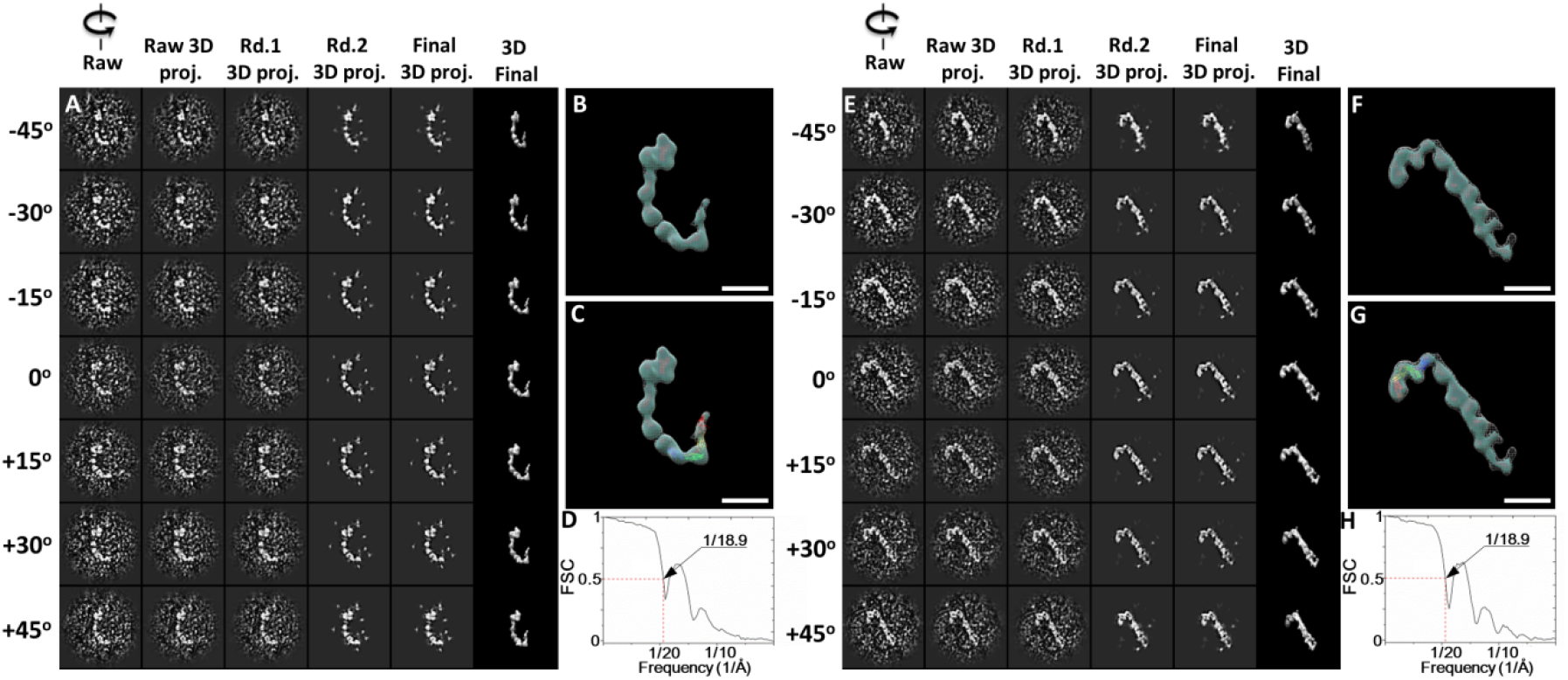
Detailed IPET 3D reconstruction process of the 21^st^ and 22^nd^ CNTN2 molecules. **(A)** Seven representative tilting views (leftmost column) of the 21^st^ individual particle were selected from 81 tilting ET micrographs after CTF correction. The corresponding tilting projections of the 3D reconstruction from major iterations are displayed beside the raw images in next four columns. Final 3D reconstruction at corresponding tilting angles is displayed in the rightmost column. **(B)** Final IPET 3D density map of the targeted individual particle displayed using two contour levels, *i.e*., 0.528 and 0.293. **(C)** Final 3D density map with the docked Ig1-Ig4 crystal structure. **(D)** The resolution of the IPET 3D map is ~18.9 Å according to FSC analyses. **(E-H)** The 3D density map of the 22^nd^ individual particle of CNTN2 was reconstructed from the tilt images using IPET. Final 3D density map was displayed using two contour levels, 0.652 and 0.210. The resolution of the IPET 3D map is ~18.9 Å according to FSC analyses. Scale bar = 100 Å.

**Figure S10.**
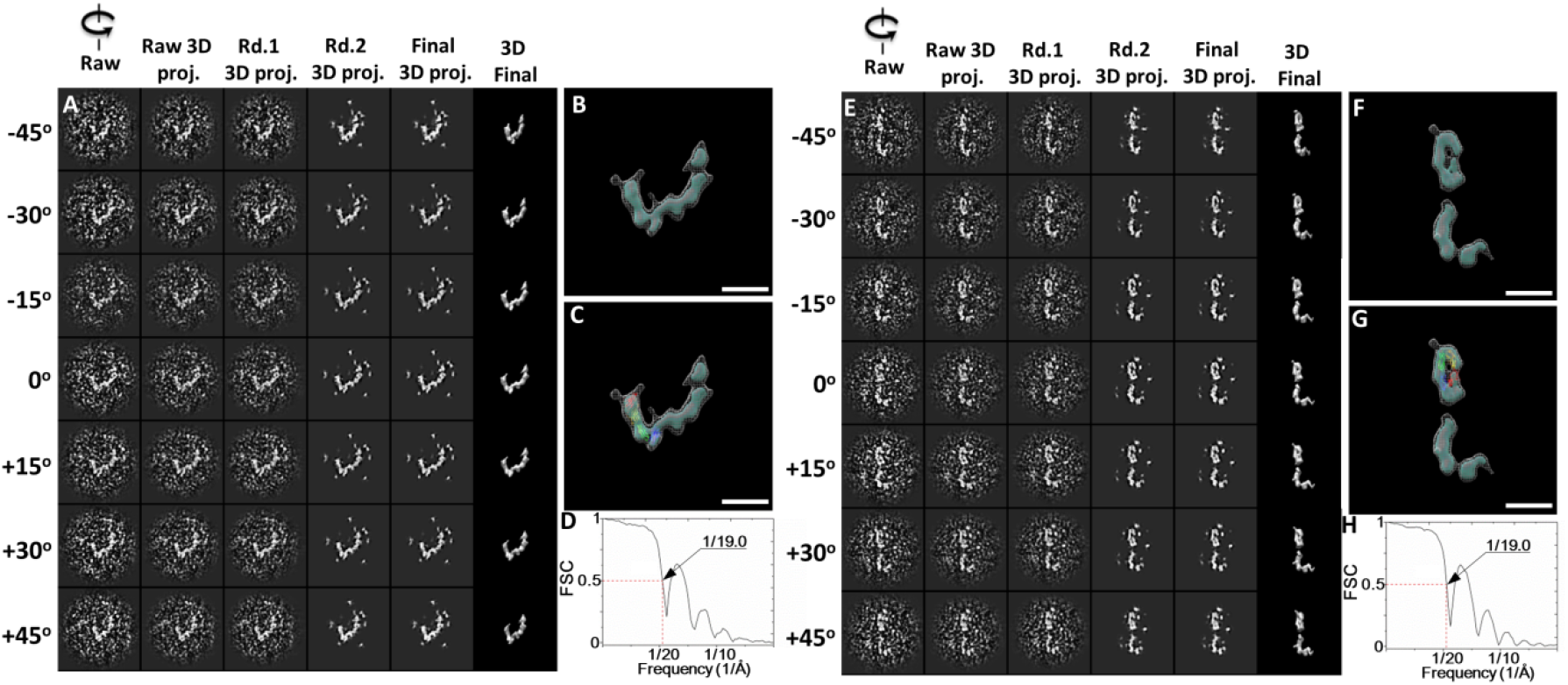
Detailed IPET 3D reconstruction process of the 23^rd^ and 24^th^ CNTN2 molecules. **(A)** Seven representative tilting views (leftmost column) of the 23^rd^ individual particle were selected from 81 tilting ET micrographs after CTF correction. The corresponding tilting projections of the 3D reconstruction from major iterations are displayed beside the raw images in next four columns. Final 3D reconstruction at corresponding tilting angles is displayed in the rightmost column. **(B)** Final IPET 3D density map of the targeted individual particle displayed using two contour levels, *i.e*., 0.617 and 0.244. **(C)** Final 3D density map with the docked Ig1-Ig4 crystal structure. **(D)** The resolution of the IPET 3D map is ~19.0 Å according to FSC analyses. **(E-H)** The 3D density map of the 12^th^ individual particle of CNTN2 was reconstructed from the tilt images using IPET. Final 3D density map was displayed using two contour levels, 0.517 and 0.227. The resolution of the IPET 3D map is ~19.0 Å according to FSC analyses. Scale bar = 100 Å.

**Figure S11.**
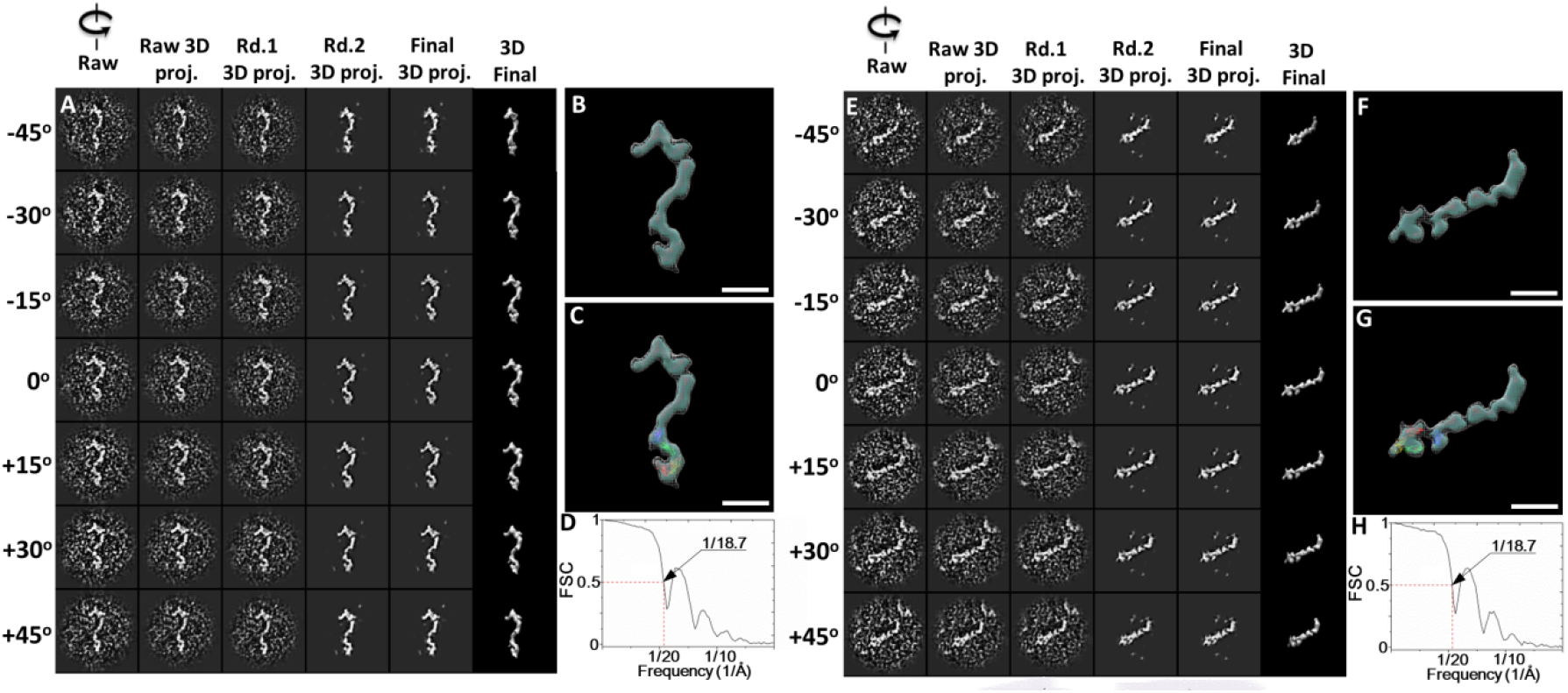
Detailed IPET 3D reconstruction process of the 25^th^ and 26^th^ CNTN2 molecules. **(A)** Seven representative tilting views (leftmost column) of the 25^th^ individual particle were selected from 81 tilting ET micrographs after CTF correction. The corresponding tilting projections of the 3D reconstruction from major iterations are displayed beside the raw images in next four columns. Final 3D reconstruction at corresponding tilting angles is displayed in the rightmost column. **(B)** Final IPET 3D density map of the targeted individual particle displayed using two contour levels, *i.e*., 0.476 and 0.335. **(C)** Final 3D density map with the docked Ig1-Ig4 crystal structure. **(D)** The resolution of the IPET 3D map is ~18.7 Å according to FSC analyses. **(E-H)** The 3D density map of the 26^th^ individual particle of CNTN2 was reconstructed from the tilt images using IPET. Final 3D density map was displayed using two contour levels, 0.502 and 0.330. The resolution of the IPET 3D map is ~18.7 Å according to FSC analyses. Scale bar = 100 Å.

**Figure S12.**
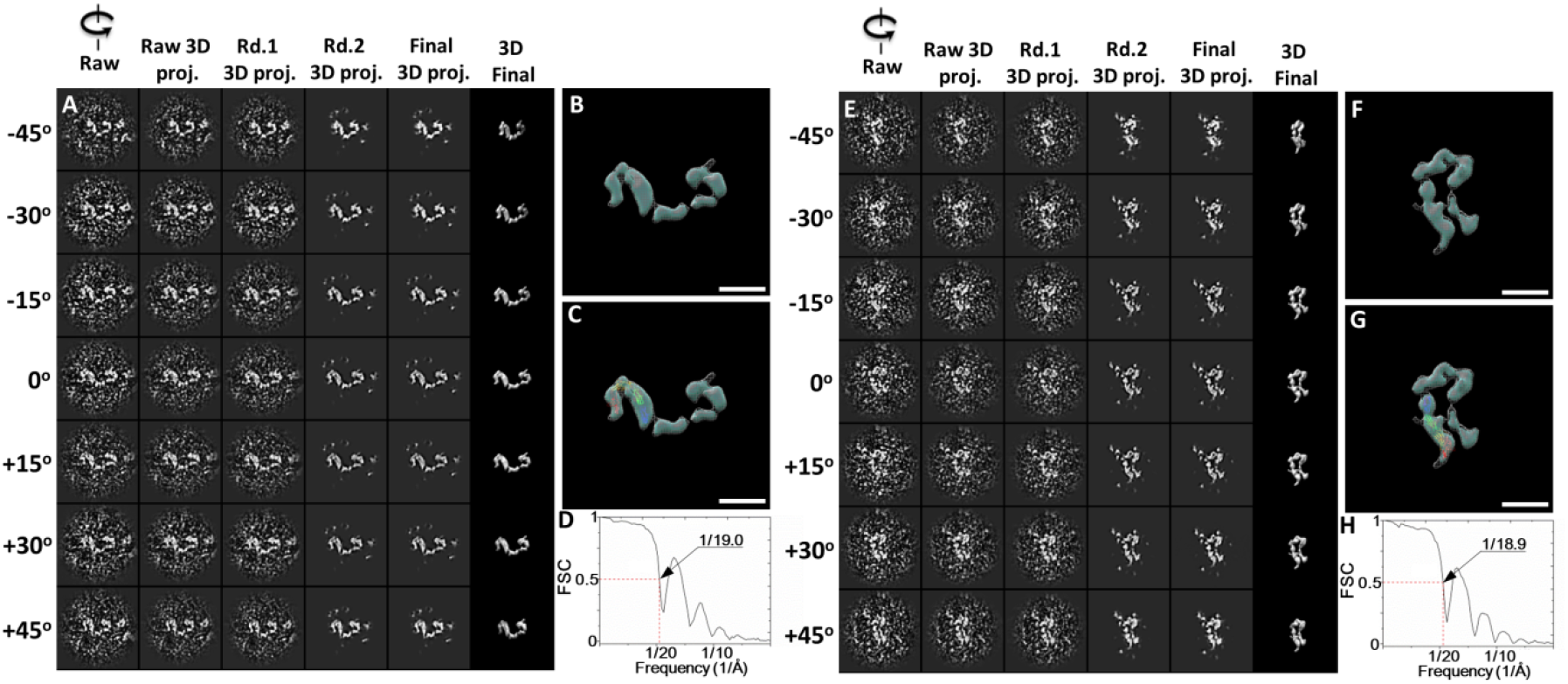
Detailed IPET 3D reconstruction process of the 27^t^ and 28^t^ CNTN2 molecules. **(A)** Seven representative tilting views (leftmost column) of the 27^th^ individual particle were selected from 81 tilting ET micrographs after CTF correction. The corresponding tilting projections of the 3D reconstruction from major iterations are displayed beside the raw images in next four columns. Final 3D reconstruction at corresponding tilting angles is displayed in the rightmost column. **(B)** Final IPET 3D density map of the targeted individual particle displayed using two contour levels, *i.e*., 0.517 and 0.251. **(C)** Final 3D density map with the docked Ig1-Ig4 crystal structure. **(D)** The resolution of the IPET 3D map is ~19.0 Å according to FSC analyses. **(E-H)** The 3D density map of the 28^th^ individual particle of CNTN2 was reconstructed from the tilt images using IPET. Final 3D density map was displayed using two contour levels, 0.459 and 0.237. The resolution of the IPET 3D map is ~18.9 Å according to FSC analyses. Scale bar = 100 Å.

**Figure S13.**
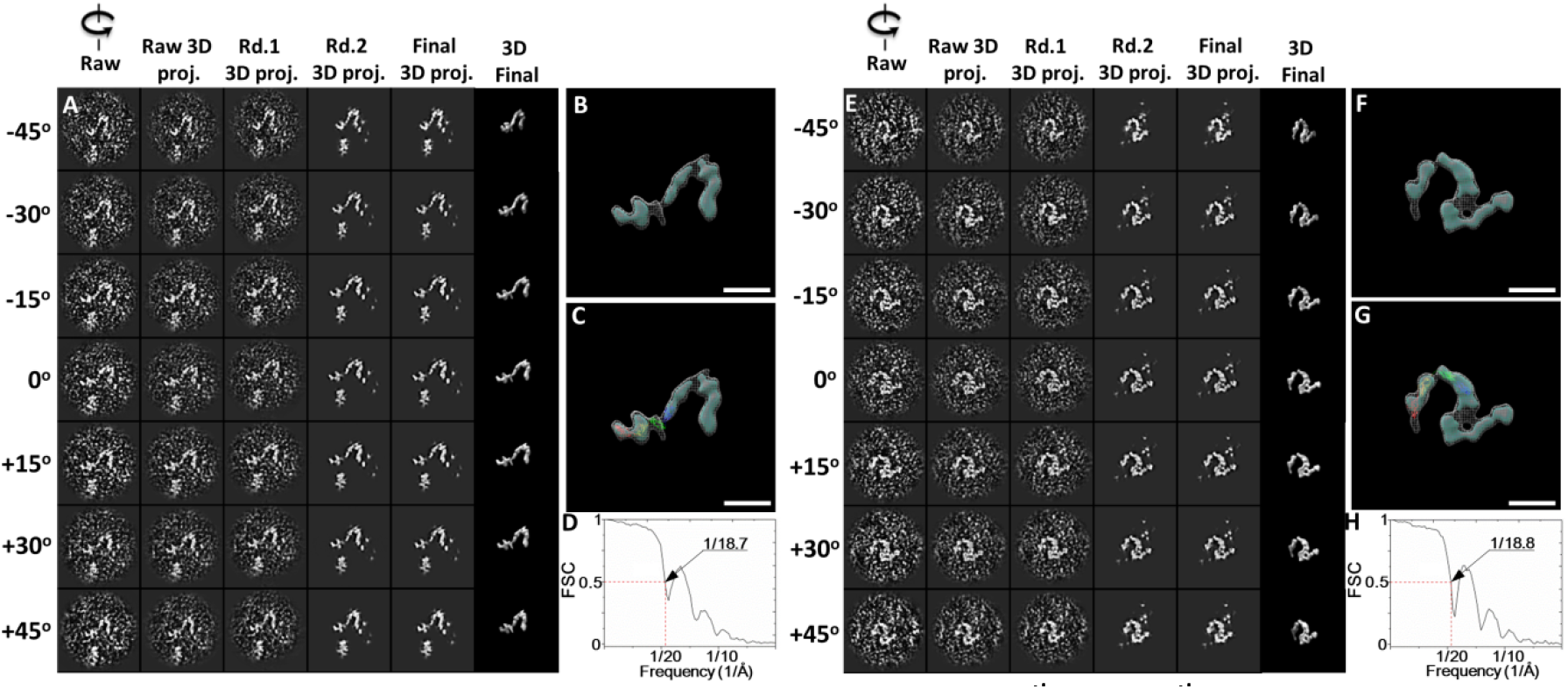
Detailed IPET 3D reconstruction process of the 29^th^ and 30^th^ CNTN2 molecules. **(A)** Seven representative tilting views (leftmost column) of the 29^th^ individual particle were selected from 81 tilting ET micrographs after CTF correction. The corresponding tilting projections of the 3D reconstruction from major iterations are displayed beside the raw images in next four columns. Final 3D reconstruction at corresponding tilting angles is displayed in the rightmost column. **(B)** Final IPET 3D density map of the targeted individual particle displayed using two contour levels, *i.e*., 0.647 and 0.261. **(C)** Final 3D density map with the docked Ig1-Ig4 crystal structure. **(D)** The resolution of the IPET 3D map is ~18.7 Å according to FSC analyses. **(E-H)** The 3D density map of the 30^th^ individual particle of CNTN2 was reconstructed from the tilt images using IPET. Final 3D density map was displayed using two contour levels, 0.542 and 0.279. The resolution of the IPET 3D map is ~18.8 Å according to FSC analyses. Scale bar = 100 Å.

**Figure S14.**
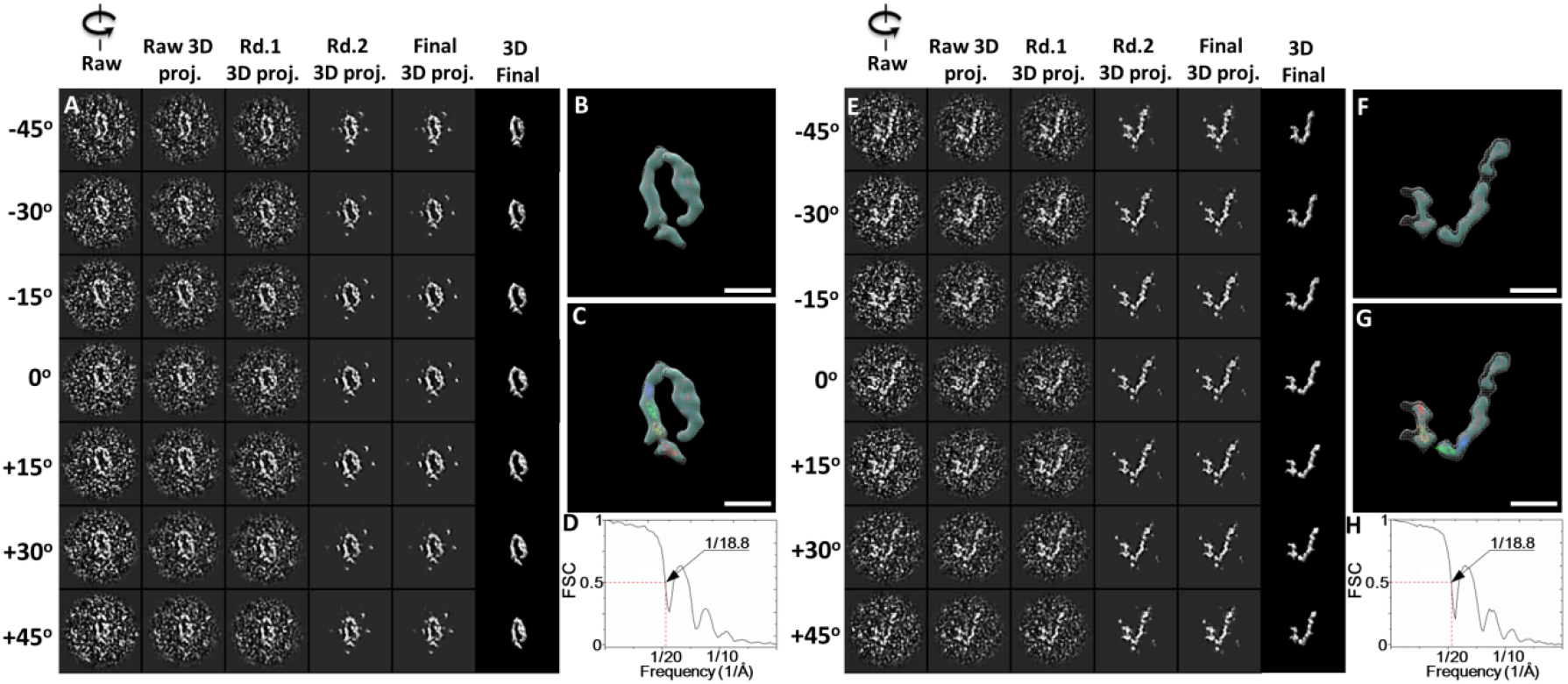
Detailed IPET 3D reconstruction process of the 31^st^ and 32^nd^ CNTN2 molecules. **(A)** Seven representative tilting views (leftmost column) of the 31^st^ individual particle were selected from 81 tilting ET micrographs after CTF correction. The corresponding tilting projections of the 3D reconstruction from major iterations are displayed beside the raw images in next four columns. Final 3D reconstruction at corresponding tilting angles is displayed in the rightmost column. **(B)** Final IPET 3D density map of the targeted individual particle displayed using two contour levels, *i.e*., 0.377 and 0.265. **(C)** Final 3D density map with the docked Ig1-Ig4 crystal structure. **(D)** The resolution of the IPET 3D map is ~18.8 Å according to FSC analyses. **(E-H)** The 3D density map of the 32^nd^ individual particle of CNTN2 was reconstructed from the tilt images using IPET. Final 3D density map was displayed using two contour levels, 0.702 and 0.328. The resolution of the IPET 3D map is ~18.8 Å according to FSC analyses. Scale bar = 100 Å.

**Figure S15.**
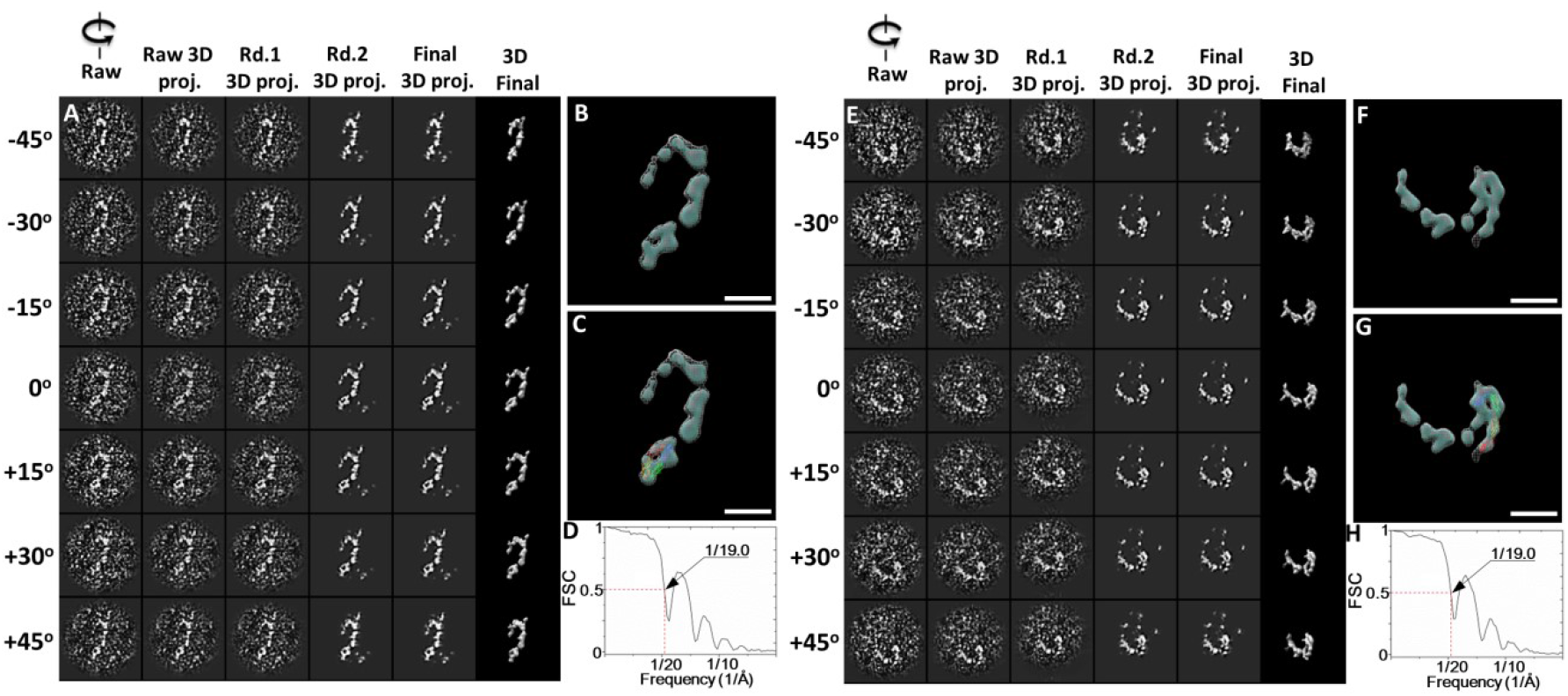
Detailed IPET 3D reconstruction process of the 33^rd^ and 34^th^ CNTN2 molecules. **(A)** Seven representative tilting views (leftmost column) of the 33^rd^ individual particle were selected from 81 tilting ET micrographs after CTF correction. The corresponding tilting projections of the 3D reconstruction from major iterations are displayed beside the raw images in next four columns. Final 3D reconstruction at corresponding tilting angles is displayed in the rightmost column. **(B)** Final IPET 3D density map of the targeted individual particle displayed using two contour levels, *i.e*., 0.516 and 0.245. **(C)** Final 3D density map with the docked Ig1-Ig4 crystal structure. **(D)** The resolution of the IPET 3D map is ~19.0 Å according to FSC analyses. **(E-H)** The 3D density map of the 34^th^ individual particle of CNTN2 was reconstructed from the tilt images using IPET. Final 3D density map was displayed using two contour levels, 0.414 and 0.296. The resolution of the IPET 3D map is ~19.0 Å according to FSC analyses. Scale bar = 100 Å.

**Figure S16.**
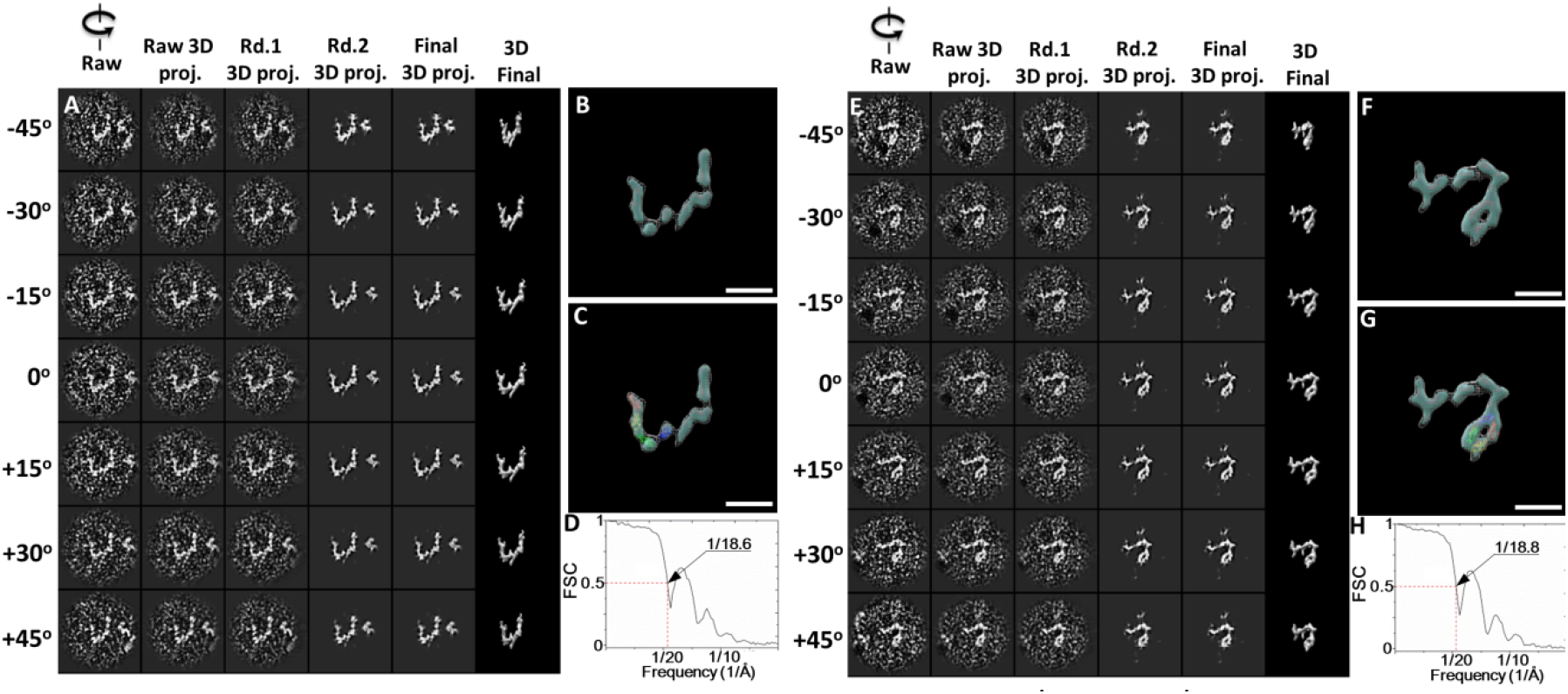
Detailed IPET 3D reconstruction process of the 35^th^ and 36^th^ CNTN2 molecules. **(A)** Seven representative tilting views (leftmost column) of the 35^th^ individual particle were selected from 81 tilting ET micrographs after CTF correction. The corresponding tilting projections of the 3D reconstruction from major iterations are displayed beside the raw images in next four columns. Final 3D reconstruction at corresponding tilting angles is displayed in the rightmost column. **(B)** Final IPET 3D density map of the targeted individual particle displayed using two contour levels, *i.e*., 0.738 and 0.505. **(C)** Final 3D density map with the docked Ig1-Ig4 crystal structure. **(D)** The resolution of the IPET 3D map is ~18.6 Å according to FSC analyses. **(E-H)** The 3D density map of the 36^th^ individual particle of CNTN2 was reconstructed from the tilt images using IPET. Final 3D density map was displayed using two contour levels, 0.550 and 0.327. The resolution of the IPET 3D map is ~18.8 Å according to FSC analyses. Scale bar = 100 Å.

**Figure S17.**
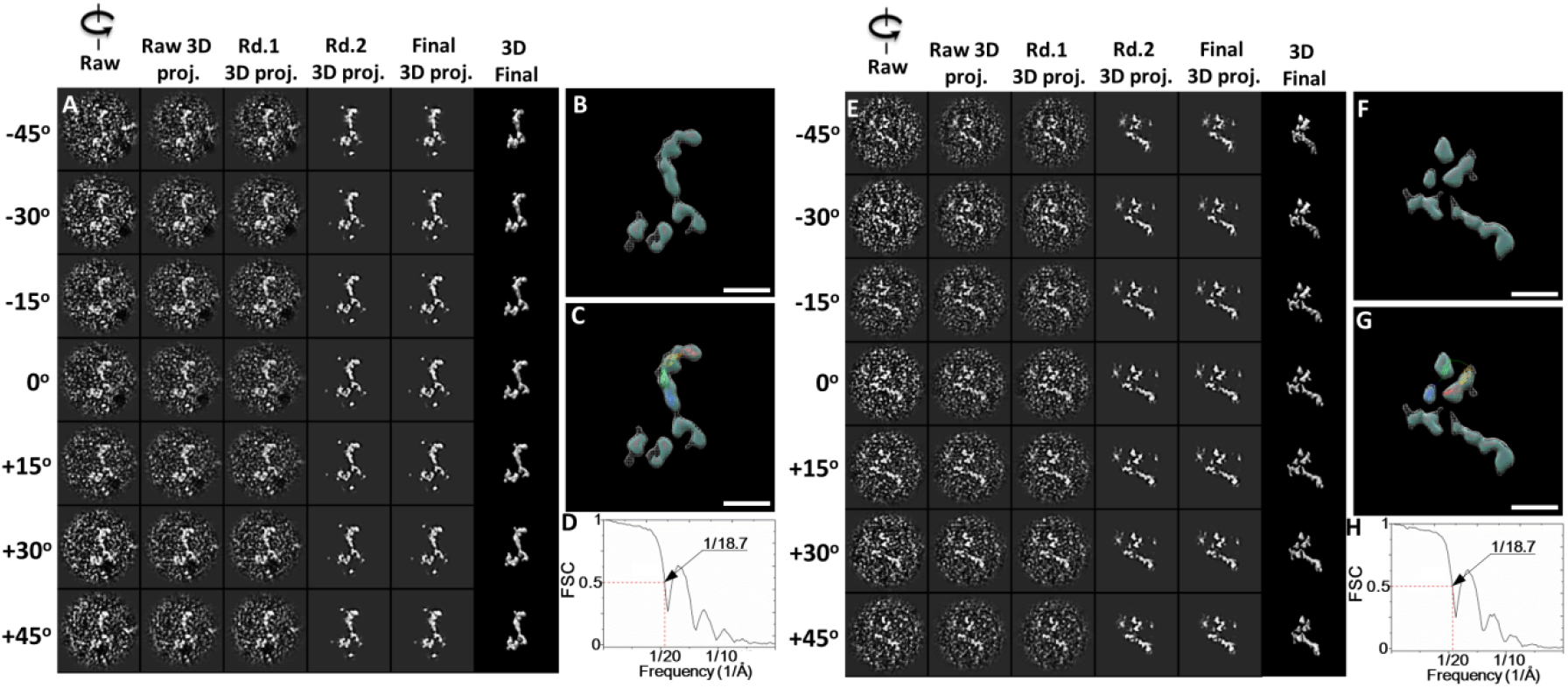
Detailed IPET 3D reconstruction process of the 37^th^ and 38^th^ CNTN2 molecules. **(A)** Seven representative tilting views (leftmost column) of the 37^th^ individual particle were selected from 81 tilting ET micrographs after CTF correction. The corresponding tilting projections of the 3D reconstruction from major iterations are displayed beside the raw images in next four columns. Final 3D reconstruction at corresponding tilting angles is displayed in the rightmost column. **(B)** Final IPET 3D density map of the targeted individual particle displayed using two contour levels, *i.e*., 0.472 and 0.283. **(C)** Final 3D density map with the docked Ig1-Ig4 crystal structure. **(D)** The resolution of the IPET 3D map is ~18.7 Å according to FSC analyses. **(E-H)** The 3D density map of the 38^th^ individual particle of CNTN2 was reconstructed from the tilt images using IPET. Final 3D density map was displayed using two contour levels, 0.292 and 0.198. The resolution of the IPET 3D map is ~18.7 Å according to FSC analyses. Scale bar = 100 Å.

**Figure S18.**
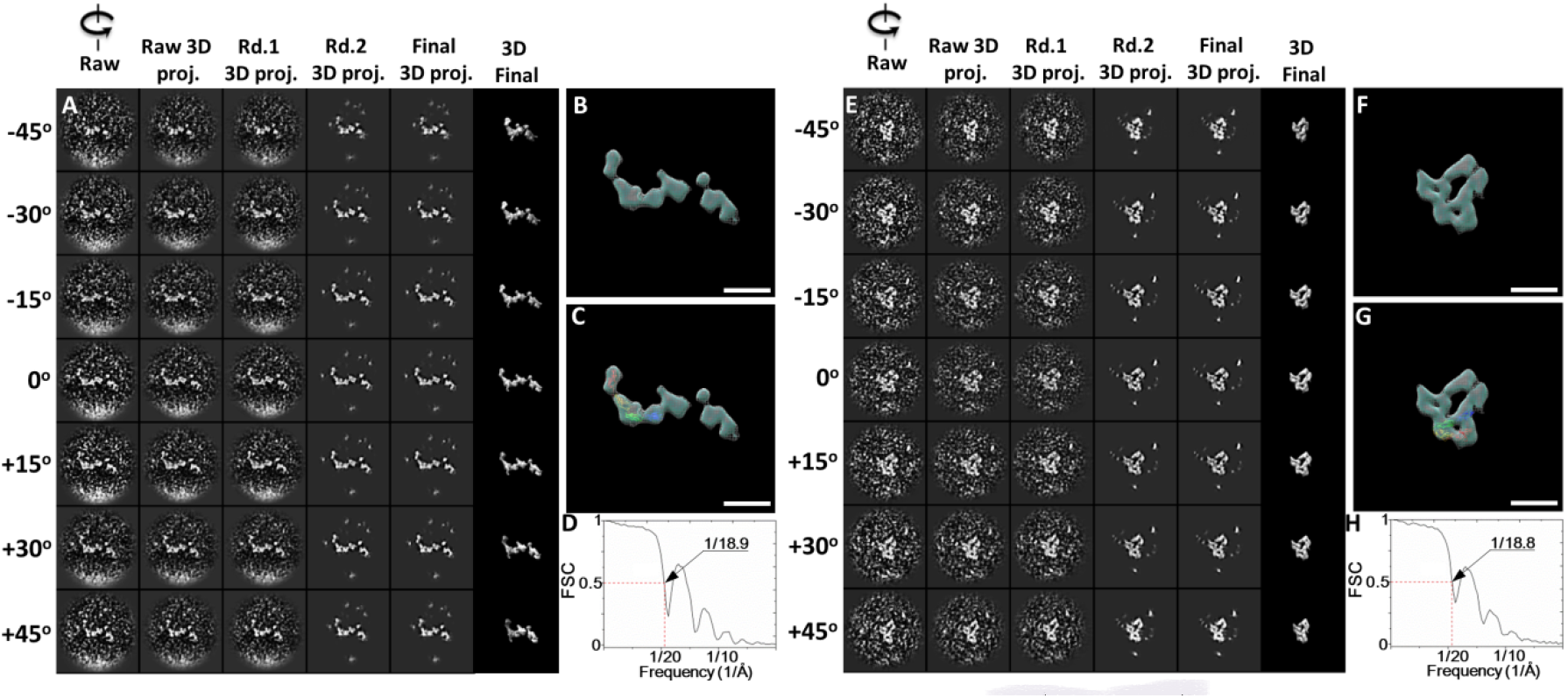
Detailed IPET 3D reconstruction process of the 39^th^ and 40^th^ CNTN2 molecules. **(A)** Seven representative tilting views (leftmost column) of the 39^th^ individual particle were selected from 81 tilting ET micrographs after CTF correction. The corresponding tilting projections of the 3D reconstruction from major iterations are displayed beside the raw images in next four columns. Final 3D reconstruction at corresponding tilting angles is displayed in the rightmost column. **(B)** Final IPET 3D density map of the targeted individual particle displayed using two contour levels, *i.e*., 0.417 and 0.305. **(C)** Final 3D density map with the docked Ig1-Ig4 crystal structure. **(D)** The resolution of the IPET 3D map is ~18.9 Å according to FSC analyses. **(E-H)** The 3D density map of the 40^th^ individual particle of CNTN2 was reconstructed from the tilt images using IPET. Final 3D density map was displayed using two contour levels, 0.530 and 0.360. The resolution of the IPET 3D map is ~18.8 Å according to FSC analyses. Scale bar = 100 Å.

**Figure S19.**
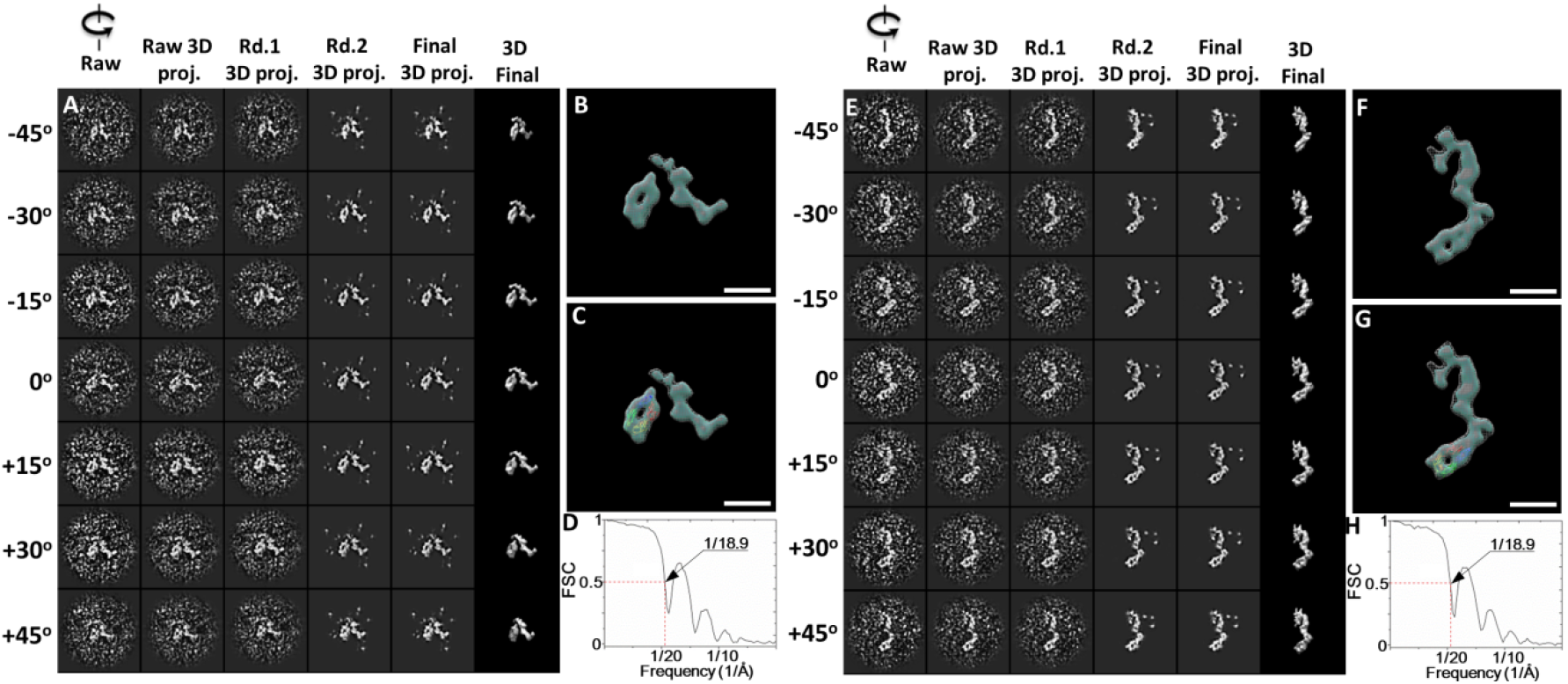
Detailed IPET 3D reconstruction process of the 41^st^ and 42^nd^ CNTN2 molecules. **(A)** Seven representative tilting views (leftmost column) of the 41^st^ individual particle were selected from 81 tilting ET micrographs after CTF correction. The corresponding tilting projections of the 3D reconstruction from major iterations are displayed beside the raw images in next four columns. Final 3D reconstruction at corresponding tilting angles is displayed in the rightmost column. **(B)** Final IPET 3D density map of the targeted individual particle displayed using two contour levels, *i.e*., 0.541 and 0.299. **(C)** Final 3D density map with the docked Ig1-Ig4 crystal structure. **(D)** The resolution of the IPET 3D map is ~18.9 Å according to FSC analyses. **(E-H)** The 3D density map of the 42^nd^ individual particle of CNTN2 was reconstructed from the tilt images using IPET. Final 3D density map was displayed using two contour levels, 0.656 and 0.339. The resolution of the IPET 3D map is ~18.9 Å according to FSC analyses. Scale bar = 100 Å.

**Figure S20.**
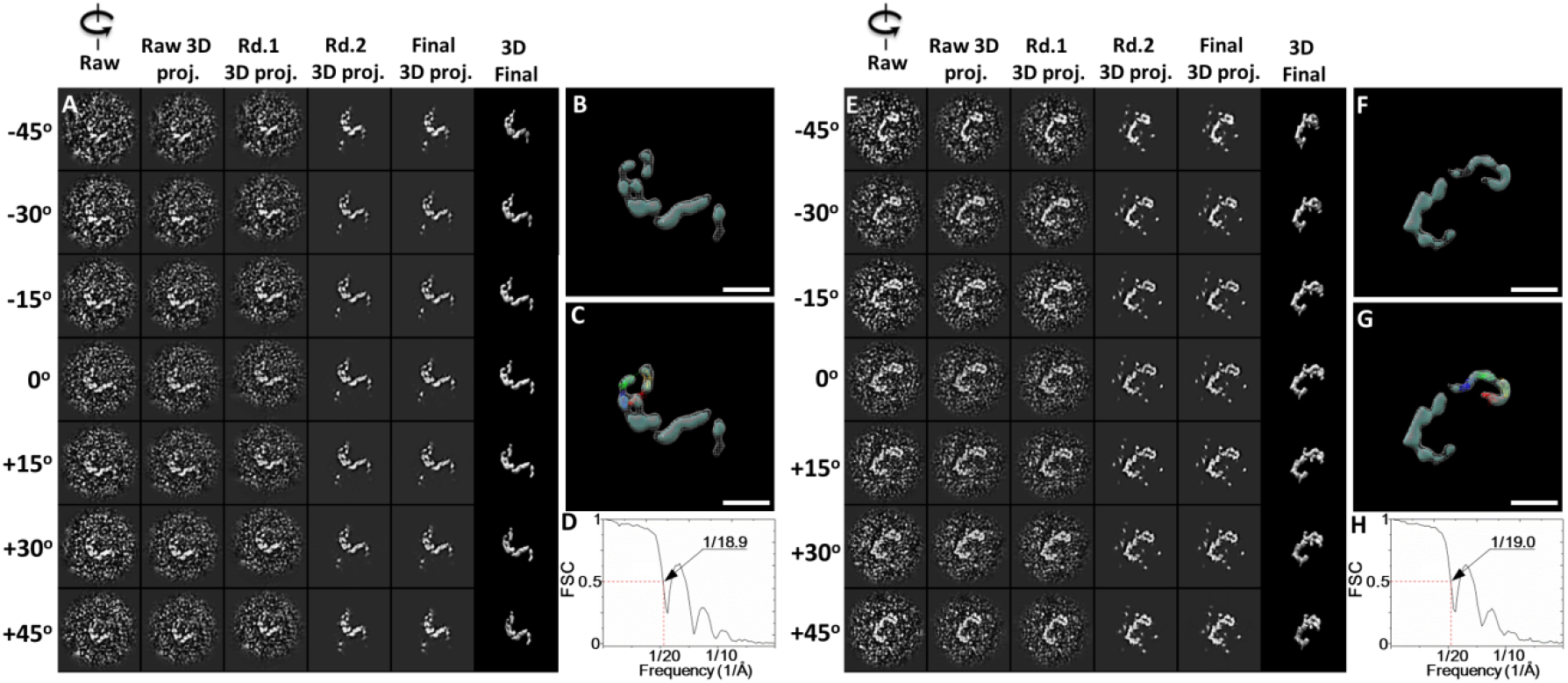
Detailed IPET 3D reconstruction process of the 43^rd^ and 44^th^ CNTN2 molecules. **(A)** Seven representative tilting views (leftmost column) of the 43^rd^ individual particle were selected from 81 tilting ET micrographs after CTF correction. The corresponding tilting projections of the 3D reconstruction from major iterations are displayed beside the raw images in next four columns. Final 3D reconstruction at corresponding tilting angles is displayed in the rightmost column. **(B)** Final IPET 3D density map of the targeted individual particle displayed using two contour levels, *i.e*., 0.687 and 0.506. **(C)** Final 3D density map with the docked Ig1-Ig4 crystal structure. **(D)** The resolution of the IPET 3D map is ~18.9 Å according to FSC analyses. **(E-H)** The 3D density map of the 44^th^ individual particle of CNTN2 was reconstructed from the tilt images using IPET. Final 3D density map was displayed using two contour levels, 0.724 and 0.310. The resolution of the IPET 3D map is ~19.0 Å according to FSC analyses. Scale bar = 100 Å.

**Figure S21.**
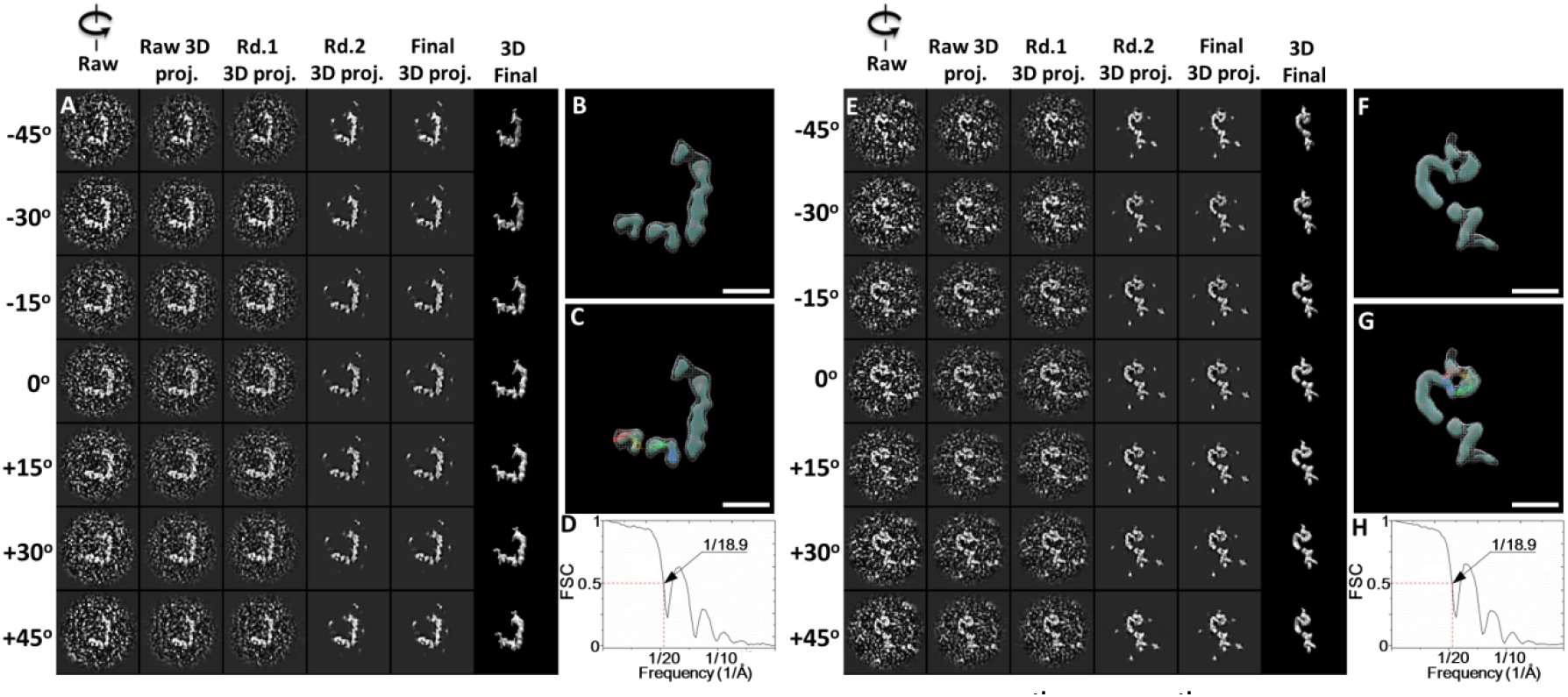
Detailed IPET 3D reconstruction process of the 45^th^ and 46^th^ CNTN2 molecules. **(A)** Seven representative tilting views (leftmost column) of the 45 ^th^ individual particle were selected from 81 tilting ET micrographs after CTF correction. The corresponding tilting projections of the 3D reconstruction from major iterations are displayed beside the raw images in next four columns. Final 3D reconstruction at corresponding tilting angles is displayed in the rightmost column. **(B)** Final IPET 3D density map of the targeted individual particle displayed using two contour levels, *i.e*., 0.520 and 0.311. **(C)** Final 3D density map with the docked Ig1-Ig4 crystal structure. **(D)** The resolution of the IPET 3D map is ~18.9 Å according to FSC analyses. **(E-H)** The 3D density map of the 46^th^ individual particle of CNTN2 was reconstructed from the tilt images using IPET. Final 3D density map was displayed using two contour levels, 0.507 and 0.262. The resolution of the IPET 3D map is ~18.9 Å according to FSC analyses. Scale bar = 100 Å.

**Figure S22.**
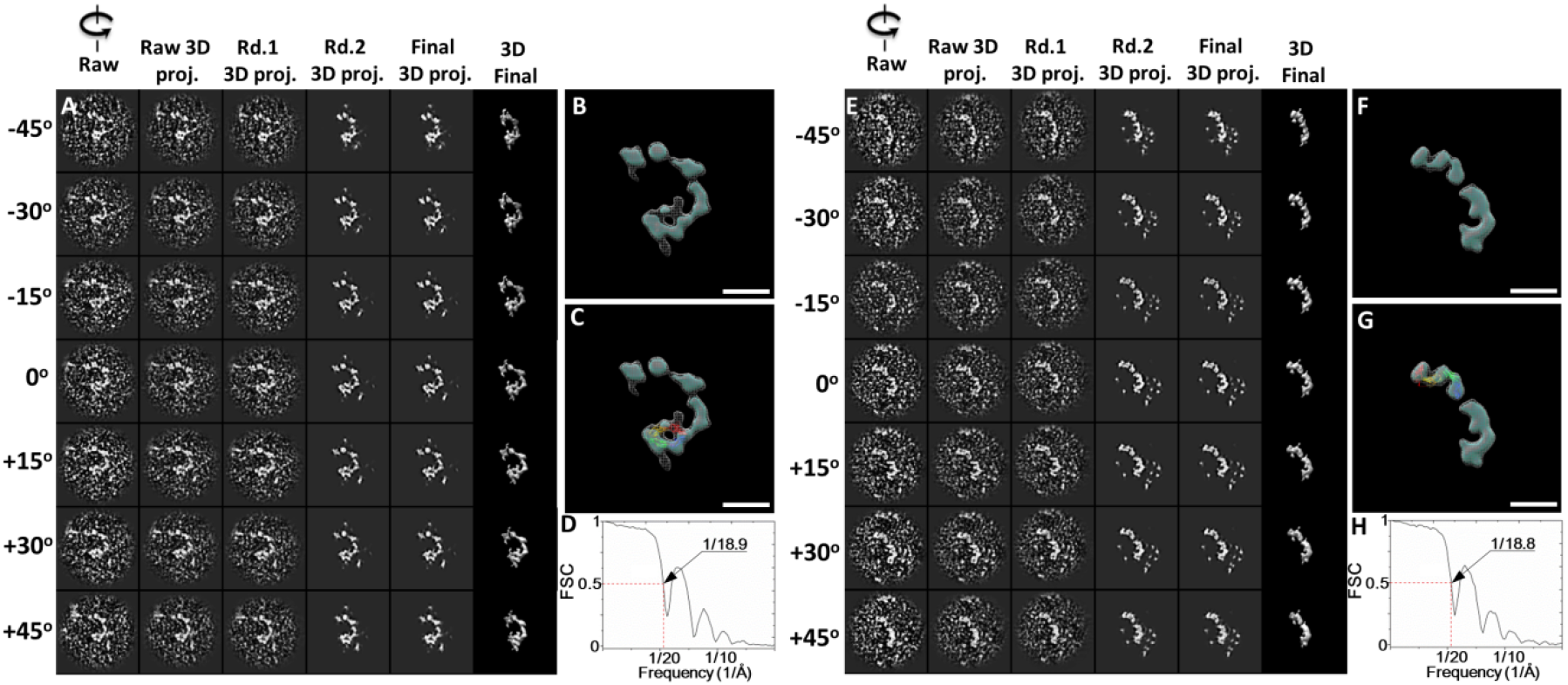
Detailed IPET 3D reconstruction process of the 47^th^ and 48^th^ CNTN2 molecules. **(A)** Seven representative tilting views (leftmost column) of the 47^th^ individual particle were selected from 81 tilting ET micrographs after CTF correction. The corresponding tilting projections of the 3D reconstruction from major iterations are displayed beside the raw images in next four columns. Final 3D reconstruction at corresponding tilting angles is displayed in the rightmost column. **(B)** Final IPET 3D density map of the targeted individual particle displayed using two contour levels, *i.e*., 0.500 and 0.312. **(C)** Final 3D density map with the docked Ig1-Ig4 crystal structure. **(D)** The resolution of the IPET 3D map is ~18.9 Å according to FSC analyses. **(E-H)** The 3D density map of the 48^th^ individual particle of CNTN2 was reconstructed from the tilt images using IPET. Final 3D density map was displayed using two contour levels, 0.786 and 0.241. The resolution of the IPET 3D map is ~18.8 Å according to FSC analyses. Scale bar = 100 Å.

**Figure S23.**
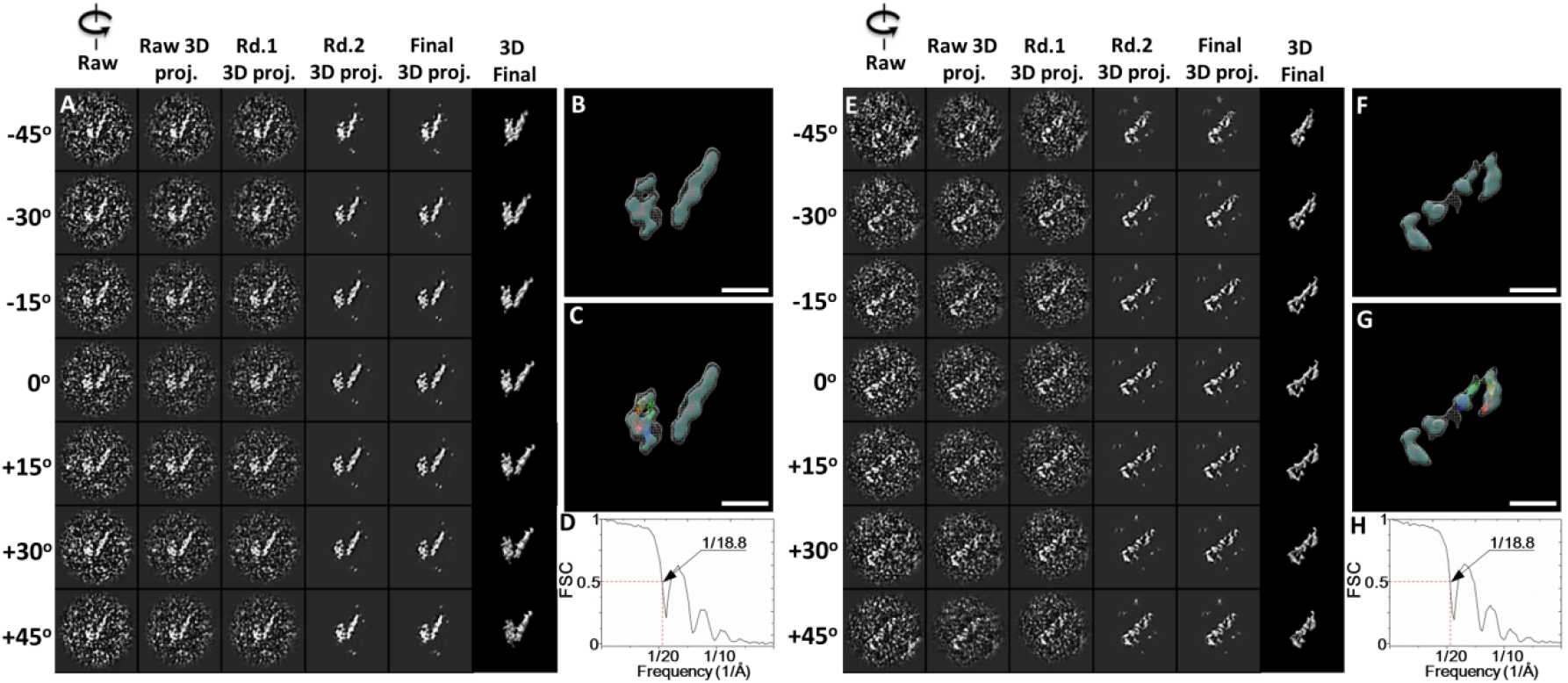
Detailed IPET 3D reconstruction process of the 49^th^ and 50^th^ CNTN2 molecules. **(A)** Seven representative tilting views (leftmost column) of the 49^th^ individual particle were selected from 81 tilting ET micrographs after CTF correction. The corresponding tilting projections of the 3D reconstruction from major iterations are displayed beside the raw images in next four columns. Final 3D reconstruction at corresponding tilting angles is displayed in the rightmost column. **(B)** Final IPET 3D density map of the targeted individual particle displayed using two contour levels, *i.e*., 0.623 and 0.278. **(C)** Final 3D density map with the docked Ig1-Ig4 crystal structure. **(D)** The resolution of the IPET 3D map is ~18.8 Å according to FSC analyses. **(E-H)** The 3D density map of the 50^th^ individual particle of CNTN2 was reconstructed from the tilt images using IPET. Final 3D density map was displayed using two contour levels, 0.574 and 0.300. The resolution of the IPET 3D map is ~18.8 Å according to FSC analyses. Scale bar = 100 Å.

**Figure S24.**
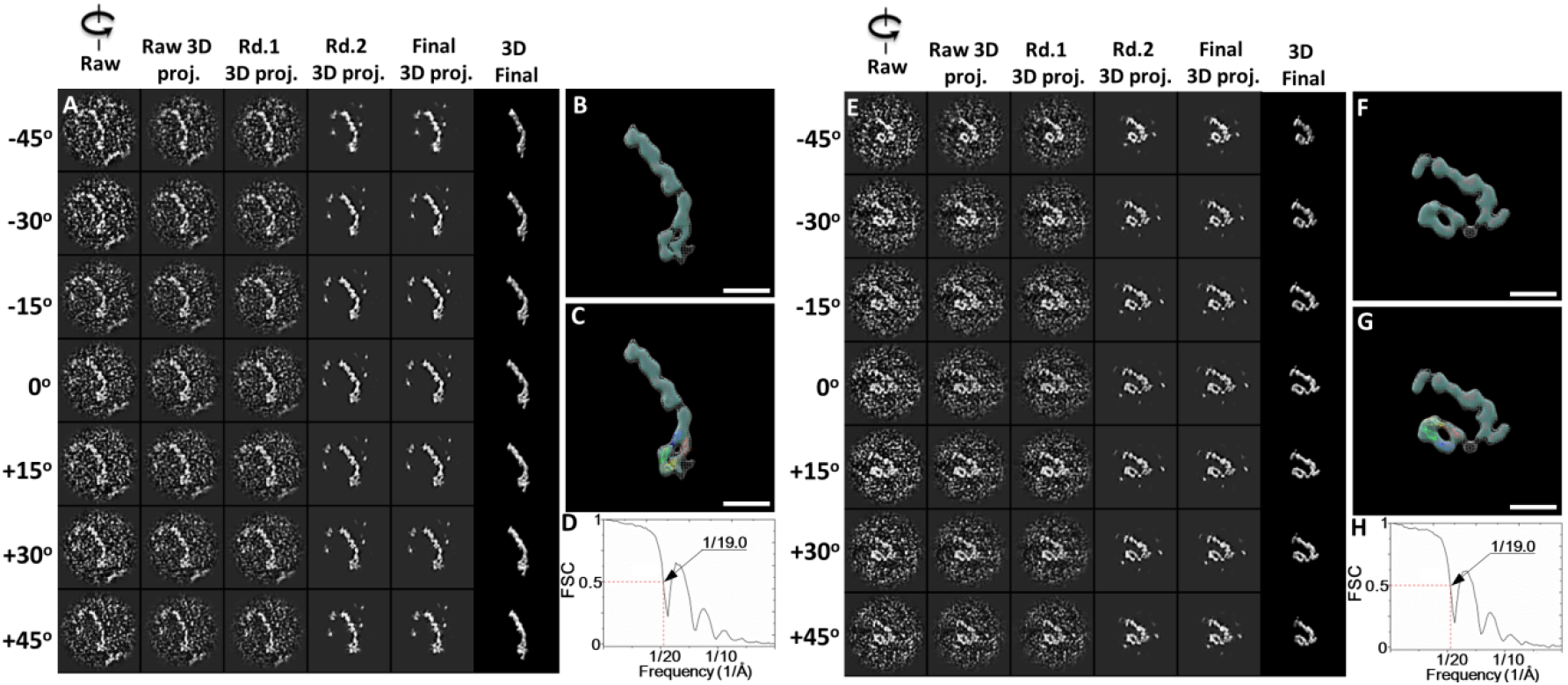
Detailed IPET 3D reconstruction process of the 51^st^ and 52^nd^ CNTN2 molecules. **(A)** Seven representative tilting views (leftmost column) of the 51^st^ individual particle were selected from 81 tilting ET micrographs after CTF correction. The corresponding tilting projections of the 3D reconstruction from major iterations are displayed beside the raw images in next four columns. Final 3D reconstruction at corresponding tilting angles is displayed in the rightmost column. **(B)** Final IPET 3D density map of the targeted individual particle displayed using two contour levels, *i.e*., 0.503 and 0.345. **(C)** Final 3D density map with the docked Ig1-Ig4 crystal structure. **(D)** The resolution of the IPET 3D map is ~19.0 Å according to FSC analyses. **(E-H)** The 3D density map of the 52^nd^ individual particle of CNTN2 was reconstructed from the tilt images using IPET. Final 3D density map was displayed using two contour levels, 0.584 and 0.167. The resolution of the IPET 3D map is ~19.0 Å according to FSC analyses. Scale bar = 100 Å.

**Figure S25.**
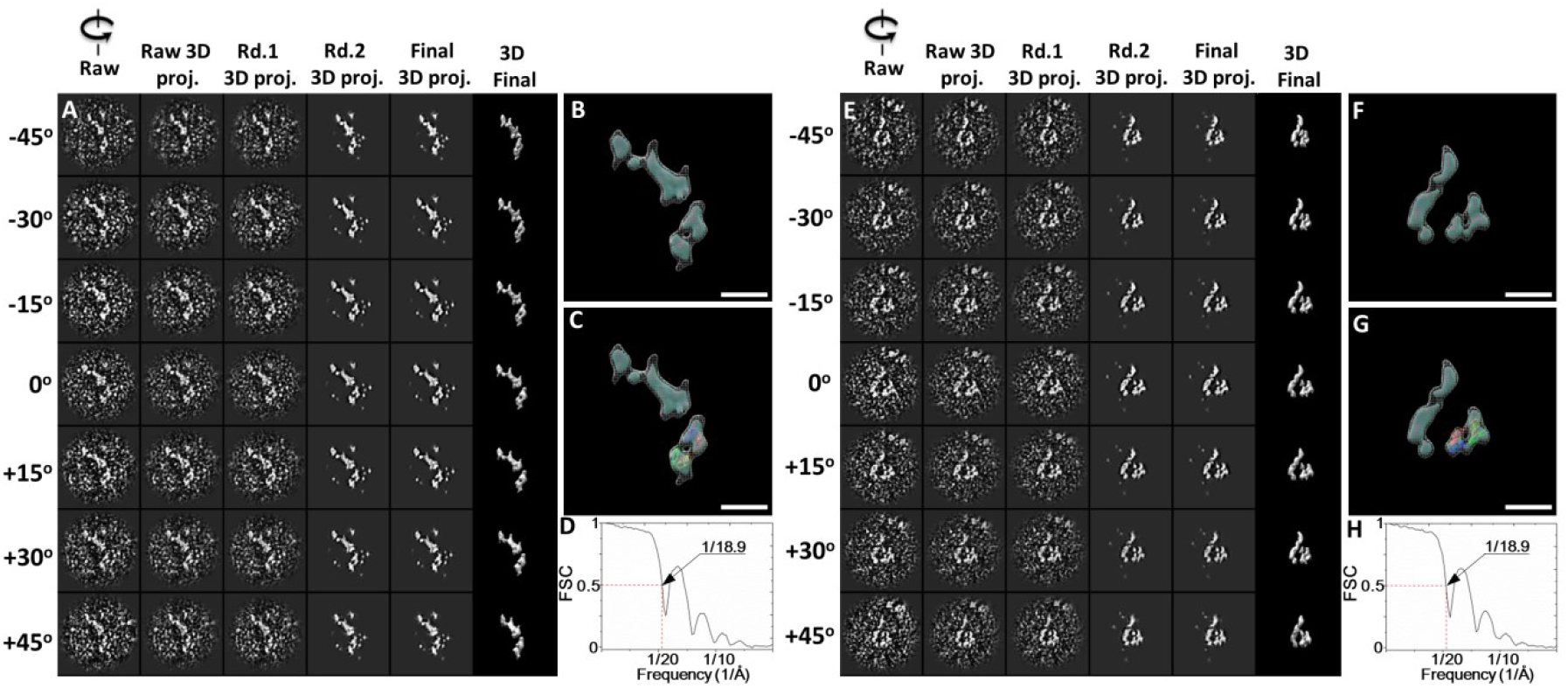
Detailed IPET 3D reconstruction process of the 53^rd^ and 54^th^ CNTN2 molecules. **(A)** Seven representative tilting views (leftmost column) of the 53^rd^ individual particle were selected from 81 tilting ET micrographs after CTF correction. The corresponding tilting projections of the 3D reconstruction from major iterations are displayed beside the raw images in next four columns. Final 3D reconstruction at corresponding tilting angles is displayed in the rightmost column. **(B)** Final IPET 3D density map of the targeted individual particle displayed using two contour levels, *i.e*., 0.689 and 0.287. **(C)** Final 3D density map with the docked Ig1-Ig4 crystal structure. **(D)** The resolution of the IPET 3D map is ~18.9 Å according to FSC analyses. **(E-H)** The 3D density map of the 54^th^ individual particle of CNTN2 was reconstructed from the tilt images using IPET. Final 3D density map was displayed using two contour levels, 0.529 and 0.279. The resolution of the IPET 3D map is ~18.9 Å according to FSC analyses. Scale bar = 100 Å.

**Figure S26.**
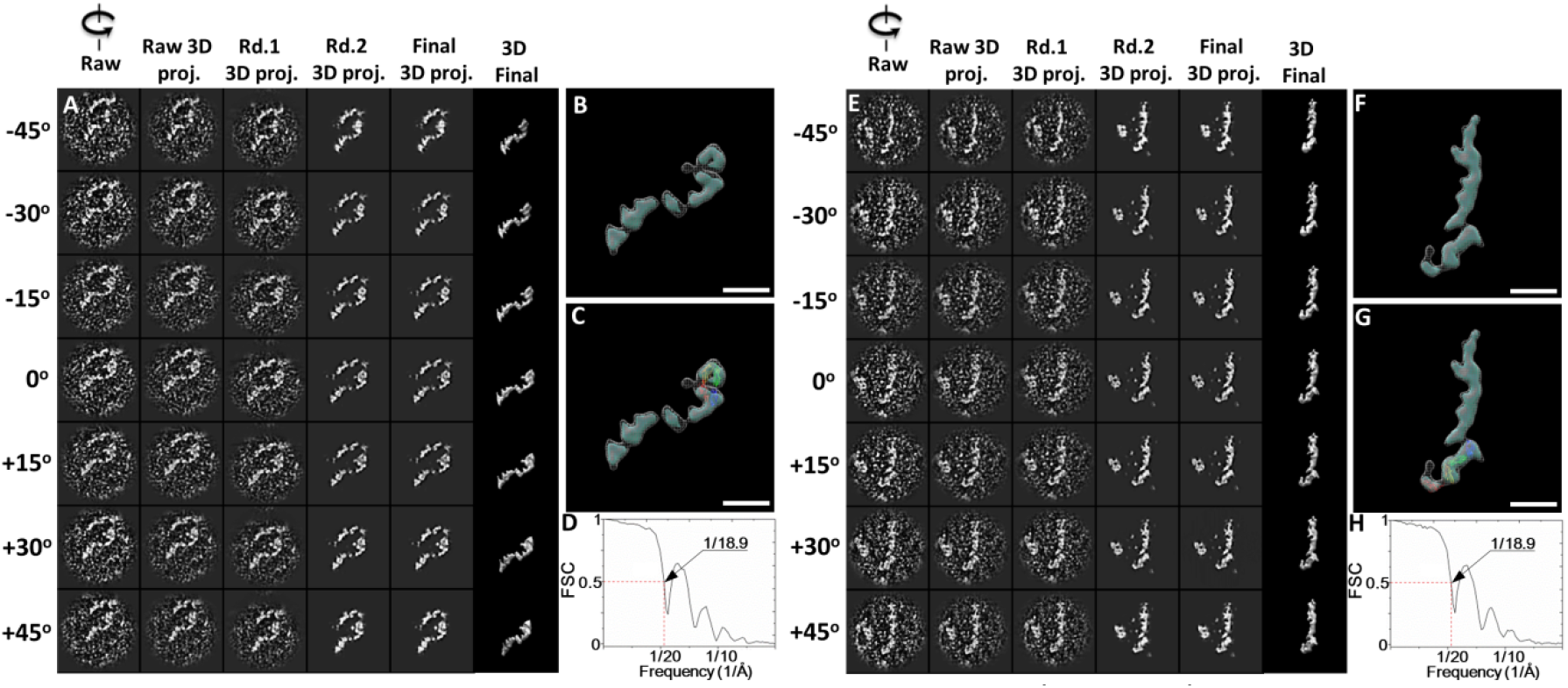
Detailed IPET 3D reconstruction process of the 55^th^ and 56^th^ CNTN2 molecules. **(A)** Seven representative tilting views (leftmost column) of the 55^th^ individual particle were selected from 81 tilting ET micrographs after CTF correction. The corresponding tilting projections of the 3D reconstruction from major iterations are displayed beside the raw images in next four columns. Final 3D reconstruction at corresponding tilting angles is displayed in the rightmost column. **(B)** Final IPET 3D density map of the targeted individual particle displayed using two contour levels, *i.e*., 0.511 and 0.340. **(C)** Final 3D density map with the docked Ig1-Ig4 crystal structure. **(D)** The resolution of the IPET 3D map is ~18.9 Å according to FSC analyses. **(E-H)** The 3D density map of the 56^th^ individual particle of CNTN2 was reconstructed from the tilt images using IPET. Final 3D density map was displayed using two contour levels, 0.583 and 0.315. The resolution of the IPET 3D map is ~18.9 Å according to FSC analyses. Scale bar = 100 Å.

**Figure S27.**
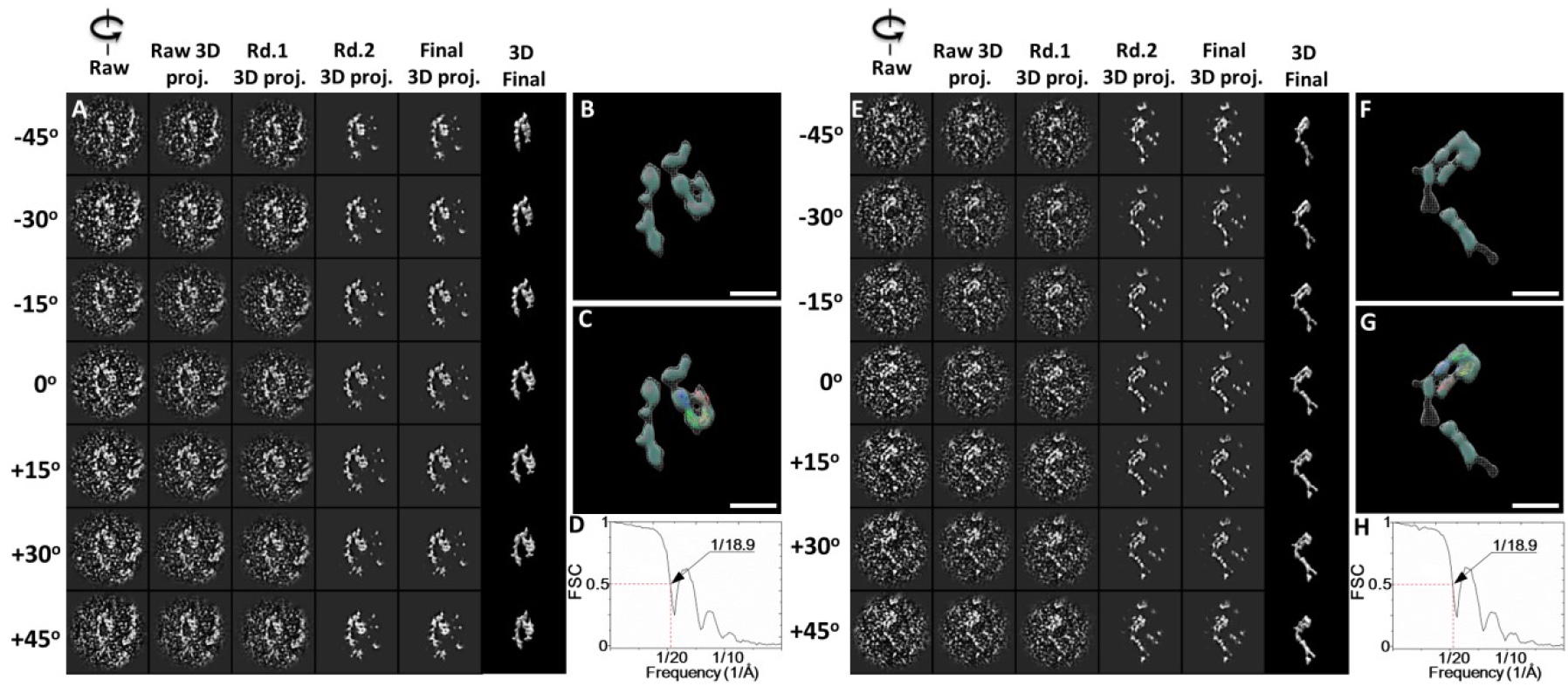
Detailed IPET 3D reconstruction process of the 57^th^ and 58^th^ CNTN2 molecules. **(A)** Seven representative tilting views (leftmost column) of the 57^th^ individual particle were selected from 81 tilting ET micrographs after CTF correction. The corresponding tilting projections of the 3D reconstruction from major iterations are displayed beside the raw images in next four columns. Final 3D reconstruction at corresponding tilting angles is displayed in the rightmost column. **(B)** Final IPET 3D density map of the targeted individual particle displayed using two contour levels, *i.e*., 0.471 and 0.236. **(C)** Final 3D density map with the docked Ig1-Ig4 crystal structure. **(D)** The resolution of the IPET 3D map is ~18.9 Å according to FSC analyses. **(E-H)** The 3D density map of the 58^th^ individual particle of CNTN2 was reconstructed from the tilt images using IPET. Final 3D density map was displayed using two contour levels, 0.569 and 0.395. The resolution of the IPET 3D map is ~18.9 Å according to FSC analyses. Scale bar = 100 Å.

**Figure S28.**
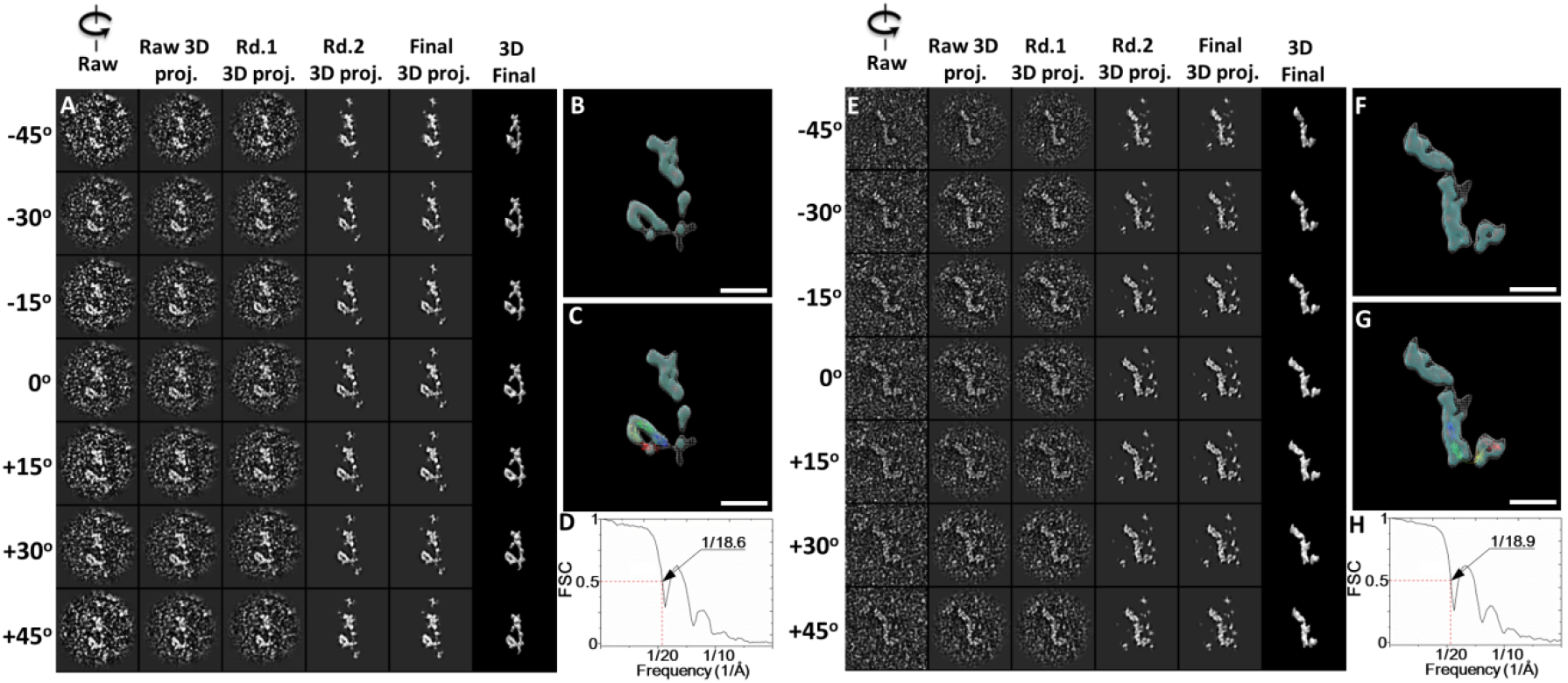
Detailed IPET 3D reconstruction process of the 59^th^ and 60^th^ CNTN2 molecules. **(A)** Seven representative tilting views (leftmost column) of the 59^th^ individual particle were selected from 81 tilting ET micrographs after CTF correction. The corresponding tilting projections of the 3D reconstruction from major iterations are displayed beside the raw images in next four columns. Final 3D reconstruction at corresponding tilting angles is displayed in the rightmost column. **(B)** Final IPET 3D density map of the targeted individual particle displayed using two contour levels, *i.e*., 0.321 and 0.167. **(C)** Final 3D density map with the docked Ig1-Ig4 crystal structure. **(D)** The resolution of the IPET 3D map is ~18.6 Å according to FSC analyses. **(E-H)** The 3D density map of the 60^th^ individual particle of CNTN2 was reconstructed from the tilt images using IPET. Final 3D density map was displayed using two contour levels, 0.211 and 0.125. The resolution of the IPET 3D map is ~18.9 Å according to FSC analyses. Scale bar = 100 Å.

**Table S1.** The parameters used for IPET 3D reconstructions

| # | EMDB#<sup>1</sup> | Dose/img<sup>2</sup><br>(e<sup>-</sup>/Å<sup>2</sup>) | Dose/se<br>t<sup>3</sup><br>(e<sup>-</sup>/Å<sup>2</sup>) | Acq. angle<br>range<sup>4</sup> | Total<br>img.<sup>5</sup> | Reconst.<br>ang. range<sup>6</sup> | Cont.<sup>7</sup> | Resol.<sup>8</sup>(Å) | Fig.<sup>9</sup> |
| --- | --- | --- | --- | --- | --- | --- | --- | --- | --- |
| 1 | EMD-7353 | 70.31 | 4570.47 | -48° to +48° | 65 | -45° to 45° | 0.516/0.302 | 18.8 | 3 |
| 2 | EMD-7354 | 43.24 | 2983.37 | -51° to +51° | 69 | -45° to 45° | 0.588/0.305 | 18.9 | 3 |
| 3 | EMD-7355 | 99.42 | 6461.98 | -48° to +48° | 65 | -45° to 45° | 0.473/0.299 | 18.5 | 3 |
| 4 | EMD-7356 | 100.03 | 6501.88 | -48° to +48° | 65 | -45° to 45° | 0.706/0.228 | 18.9 | 3 |
| 5 | EMD-7357 | 98.94 | 6431.26 | -48° to +48° | 65 | -45° to 45° | 0.211/0.125 | 18.7 | S1 |
| 6 | EMD-7358 | 71.24 | 4630.30 | -48° to +48° | 65 | -45° to 45° | 0.326/0.159 | 18.9 | S1 |
| 7 | EMD-7359 | 100.13 | 6508.28 | -48° to +48° | 65 | -45° to 45° | 0.230/0.151 | 18.9 | S2 |
| 8 | EMD-7360 | 70.09 | 4556.14 | -48° to +48° | 65 | -45° to 45° | 0.572/0.342 | 18.9 | S2 |
| 9 | EMD-7361 | 70.57 | 4586.78 | -48° to +48° | 65 | -45° to 45° | 0.286/0.169 | 18.9 | S3 |
| 10 | EMD-7362 | 70.10 | 4556.44 | -48° to +48° | 65 | -45° to 45° | 0.421/0.309 | 18.9 | S3 |
| 11 | EMD-7363 | 40.72 | 2809.94 | -51° to +51° | 69 | -45° to 45° | 0.252/0.135 | 18.7 | S4 |
| 12 | EMD-7364 | 43.73 | 3017.69 | -51° to +51° | 69 | -45° to 45° | 0.267/0.154 | 18.7 | S4 |
| 13 | EMD-7365 | 41.79 | 2883.78 | -51° to +51° | 69 | -45° to 45° | 0.629/0.360 | 19.4 | S5 |
| 14 | EMD-7366 | 41.36 | 2854.06 | -51° to +51° | 69 | -45° to 45° | 0.217/0.141 | 19.4 | S5 |
| 15 | EMD-7367 | 70.30 | 4569.18 | -48° to +48° | 65 | -45° to 45° | 0.245/0.151 | 19.2 | S6 |
| 16 | EMD-7368 | 97.65 | 6347.29 | -48° to +48° | 65 | -45° to 45° | 0.529/0.177 | 19.0 | S6 |
| 17 | EMD-7369 | 70.75 | 4598.56 | -48° to +48° | 65 | -45° to 45° | 0.400/0.258 | 19.0 | S7 |
| 18 | EMD-7370 | 42.66 | 2943.60 | -51° to +51° | 69 | -45° to 45° | 0.668/0.321 | 19.0 | S7 |
| 19 | EMD-7371 | 99.26 | 6451.61 | -48° to +48° | 65 | -45° to 45° | 0.415/0.236 | 18.5 | S8 |
| 20 | EMD-7372 | 71.10 | 4621.51 | -48° to +48° | 65 | -45° to 45° | 0.355/0.274 | 18.9 | S8 |
| 21 | EMD-7373 | 43.17 | 2978.80 | -51° to +51° | 69 | -45° to 45° | 0.528/0.293 | 18.9 | S9 |
| 22 | EMD-7374 | 39.99 | 2759.35 | -51° to +51° | 69 | -45° to 45° | 0.652/0.210 | 18.9 | S9 |
| 23 | EMD-7375 | 100.03 | 6501.94 | -48° to +48° | 65 | -45° to 45° | 0.617/0.244 | 19.0 | S10 |
| 24 | EMD-7376 | 70.34 | 4571.86 | -48° to +48° | 65 | -45° to 45° | 0.517/0.227 | 19.0 | S10 |
| 25 | EMD-7377 | 70.07 | 4554.84 | -48° to +48° | 65 | -45° to 45° | 0.476/0.335 | 18.7 | S11 |
| 26 | EMD-7378 | 71.03 | 4616.89 | -48° to +48° | 65 | -45° to 45° | 0.502/0.330 | 18.7 | S11 |
| 27 | EMD-7379 | 70.50 | 4582.44 | -48° to +48° | 65 | -45° to 45° | 0.517/0.251 | 19.0 | S12 |
| 28 | EMD-7380 | 70.31 | 4570.41 | -48° to +48° | 65 | -45° to 45° | 0.459/0.237 | 18.9 | S12 |
| 29 | EMD-7381 | 41.74 | 2880.14 | -51° to +51° | 69 | -45° to 45° | 0.647/0.261 | 18.7 | S13 |
| 30 | EMD-7382 | 40.76 | 2812.50 | -51° to +51° | 69 | -45° to 45° | 0.542/0.279 | 18.8 | S13 |
| 31 | EMD-7383 | 99.09 | 6440.60 | -48° to +48° | 65 | -45° to 45° | 0.377/0.265 | 18.8 | S14 |
| 32 | EMD-7384 | 95.16 | 6185.12 | -48° to +48° | 65 | -45° to 45° | 0.702/0.328 | 18.8 | S14 |
| 33 | EMD-7385 | 100.38 | 6524.42 | -48° to +48° | 65 | -45° to 45° | 0.516/0.245 | 19.0 | S15 |
| 34 | EMD-7386 | 71.09 | 4620.67 | -48° to +48° | 65 | -45° to 45° | 0.414/0.296 | 19.0 | S15 |
| 35 | EMD-7387 | 44.02 | 3037.25 | -51° to +51° | 69 | -45° to 45° | 0.738/0.505 | 18.6 | S16 |
| 36 | EMD-7388 | 70.48 | 4581.01 | -48° to +48° | 65 | -45° to 45° | 0.550/0.327 | 18.8 | S16 |
| 37 | EMD-7389 | 70.14 | 4558.92 | -48° to +48° | 65 | -45° to 45° | 0.472/0.283 | 18.7 | S17 |
| 38 | EMD-7390 | 70.15 | 4559.58 | -48° to +48° | 65 | -45° to 45° | 0.292/0.198 | 18.7 | S17 |
| 39 | EMD-7391 | 70.63 | 4591.20 | -48° to +48° | 65 | -45° to 45° | 0.417/0.305 | 18.9 | S18 |
| 40 | EMD-7392 | 99.79 | 6486.67 | -48° to +48° | 65 | -45° to 45° | 0.530/0.360 | 18.8 | S18 |
| 41 | EMD-7393 | 70.44 | 4578.89 | -48° to +48° | 65 | -45° to 45° | 0.541/0.299 | 18.9 | S19 |
| 42 | EMD-7394 | 70.68 | 4593.88 | -48° to +48° | 65 | -45° to 45° | 0.656/0.339 | 18.9 | S19 |

| 43 | EMD-7395 | 44.29 | 3056.08 | -51° to +51° | 69 | -45° to 45° | 0.687/0.506 | 18.9 | S20 |
| --- | --- | --- | --- | --- | --- | --- | --- | --- | --- |
| 44 | EMD-7396 | 43.16 | 2978.33 | -51° to +51° | 69 | -45° to 45° | 0.724/0.310 | 19.0 | S20 |
| 45 | EMD-7397 | 100.25 | 6516.05 | -48° to +48° | 65 | -45° to 45° | 0.520/0.311 | 18.9 | S21 |
| 46 | EMD-7398 | 70.44 | 4578.57 | -48° to +48° | 65 | -45° to 45° | 0.507/0.262 | 18.9 | S21 |
| 47 | EMD-7399 | 70.57 | 4587.12 | -48° to +48° | 65 | -45° to 45° | 0.500/0.312 | 18.9 | S22 |
| 48 | EMD-7400 | 40.53 | 2796.79 | -51° to +51° | 69 | -45° to 45° | 0.786/0.241 | 18.8 | S22 |
| 49 | EMD-7401 | 70.63 | 4590.67 | -48° to +48° | 65 | -45° to 45° | 0.623/0.278 | 18.8 | S23 |
| 50 | EMD-7402 | 70.23 | 4564.90 | -48° to +48° | 65 | -45° to 45° | 0.574/0.300 | 18.8 | S23 |
| 51 | EMD-7404 | 43.85 | 3025.72 | -51° to +51° | 69 | -45° to 45° | 0.503/0.345 | 19.0 | S24 |
| 52 | EMD-7405 | 70.63 | 4590.93 | -48° to +48° | 65 | -45° to 45° | 0.584/0.167 | 19.0 | S24 |
| 53 | EMD-7406 | 70.70 | 4595.56 | -48° to +48° | 65 | -45° to 45° | 0.689/0.287 | 18.9 | S25 |
| 54 | EMD-7407 | 70.50 | 4582.72 | -48° to +48° | 65 | -45° to 45° | 0.529/0.279 | 18.9 | S25 |
| 55 | EMD-7408 | 70.61 | 4589.47 | -48° to +48° | 65 | -45° to 45° | 0.511/0.340 | 18.9 | S26 |
| 56 | EMD-7409 | 70.90 | 4608.70 | -48° to +48° | 65 | -45° to 45° | 0.583/0.315 | 18.9 | S26 |
| 57 | EMD-7410 | 70.12 | 4557.54 | -48° to +48° | 65 | -45° to 45° | 0.471/0.236 | 18.9 | S27 |
| 58 | EMD-7411 | 70.87 | 4606.42 | -48° to +48° | 65 | -45° to 45° | 0.569/0.395 | 18.9 | S27 |
| 59 | EMD-7412 | 69.81 | 4537.84 | -48° to +48° | 65 | -45° to 45° | 0.321/0.167 | 18.6 | S28 |
| 60 | EMD-7413 | 69.72 | 4531.56 | -48° to +48° | 65 | -45° to 45° | 0.211/0.125 | 18.9 | S28 |
| # | EMDB#<sup>1</sup> | Dose/img<sup>2</sup><br>(e<sup>-</sup>/Å<sup>2</sup>) | Dose/se<sup>3</sup><br>(e<sup>-</sup>/Å<sup>2</sup>) | Acq. angle<br>range<sup>4</sup> | Total<br>img.<sup>5</sup> | Reconst.<br>ang. Range<sup>6</sup> | Cont.<sup>7</sup> | Resol.<sup>8</sup><br>(Å) | Fig.<sup>9</sup> |

<sup>1</sup> EMDB Index: https://www.ebi.ac.uk/pdbe/emdb/
<sup>2</sup> Dose used for each CCD frame
<sup>3</sup> Dose used for whole tilt series
<sup>4</sup> Data acquisition angle range
<sup>5</sup> Total images in the tilt series
<sup>6</sup> Reconstruction angle range
<sup>7</sup> Contour used for display
<sup>8</sup> IPET 3D reconstruction resolution
<sup>9</sup> The process of IPET 3D reconstruction showed in Figure and Supplementary Figure
TEM model: Zeiss 120 stands for Zeiss Libra 120 Plus TEM
CCD: UltraScan for Gatan UltraScan 4000 4Kx4K CCD
Angstrom per pixel: 2.96 Å (after bin=2)

